# Assembly of a pangenome uncovers novel non-reference unique insertion sequences in cattle highlighting their genetic diversity

**DOI:** 10.64898/2025.12.02.691810

**Authors:** Valentin Sorin, Florian Besnard, Aurélien Capitan, Cécile Grohs, Maulana Mughitz Naji, Clémentine Escouflaire, Sébastien Fritz, Joanna Lledo, Camille Eché, Carole Iampietro, Cécile Donnadieu, Denis Milan, Laurence Drouilhet, Gwenola Tosser-Klopp, Didier Boichard, Christophe Klopp, Marie-Pierre Sanchez, Mekki Boussaha

## Abstract

**Background:** The current cattle reference genome, derived from a single Hereford cow, does not capture the full spectrum of genetic diversity present within the species. Moreover, detecting structural variations (SVs ≥ 50 nucleotides long) remains challenging using only standard approaches of either short or long-read sequence approaches against a linear reference genome. Recent advances in long-read sequencing technologies and graph-based assembly now enable the construction of breed-specific pangenomes, revealing previously uncharacterized genomic regions that may contribute to important agricultural traits.

**Results:** In this study we constructed a cattle pangenome graph using 16 high-quality haplotype-resolved genome assemblies originating from nine breeds representing the diversity of French cattle populations, and including Yak (*Bos grunniens*) as a close outgroup species. Using a trio-based strategy combined with complementary sequencing technologies and bioinformatics methods, we identified and characterized 101,219 structural variations. Of these, 33,634 were classified as non-reference unique insertions (NRUIs), adding several megabases of novel genomic sequences absent from the current Hereford reference genome. Analysis of the distribution of these NRUIs revealed significant genome-wide enrichment within QTL regions associated with milk production and morphological traits, suggesting their contribution to the genetic basis of economically relevant phenotypes. Furthermore, their functional annotation highlighted two NRUIs located within the intronic regions of *ARMH3* and *EPHA5*, both specific to the Normande breed and significantly associated with milk production and morphological traits, respectively.

**Conclusions:** Our findings demonstrate the value of pangenome approaches to uncover functionally relevant SVs, particularly NRUIs, that are systematically not in the current reference genome. By linking these variants to economically important traits, our work underscores the need to incorporate breed diversity into future genomic analyses and reference-building efforts in cattle.

## Background

The advent of long-read sequencing technologies that allow for the production of sequences for longer fragments has furthered our comprehension of the bovine genome. Over the past decade, significant efforts have been undertaken to produce *de novo* genome assemblies, enabling the development of pangenomic studies in cattle.

These initiatives have resulted in the creation of the international Bovine Pangenome Consortium (https://bovinepangenome.github.io/), whose primary objective is to construct a comprehensive pangenome of domestic cattle and closely related species to identify variants lost during domestication or breed differentiation [1].

Recent studies have confirmed the accuracy and robustness of cattle pangenomes: For instance, Leonard *et al.* [2] have reported high levels of concordance in SV detection across multiple sequencing platforms; while another study revealed strong agreement among various pangenome construction methods [3]. However, these studies also pointed out that each method had particular advantages and disadvantages that should be considered when examining variants derived from multiple assemblies.

In cattle, several studies have demonstrated the potential of pangenomes to identify functionally relevant SVs. One study identified a complex structural variant located upstream of the *KIT* gene, which is strongly associated with the white-headed coat pattern in various cattle breeds [4]. Another study developed a pangenome comprising 64 *de novo* assemblies from 14 French breeds, revealing an 8.2 kb deletion within *MATN3* significantly associated with stature in Holstein, as well as a high-quality panel of non-reference unique insertions (NRUIs), totalling a size of 25 Mb [5].

The use of a single linear reference assembly limits the characterization of cattle genetic diversity, particularly for NRUIs, which represent megabases of genomic sequences that are missing from the current bovine reference genome assembly that may harbour alleles influencing economically important traits in cattle.

A pangenomic study, based on a bovine multi-assembly graph constructed with three breeds of taurine cattle (Angus, Highland, and Original Braunvieh) and two close relatives, Brahman (*Bos taurus indicus*) and yak (*B. grunniens*), uncovered 70 Mb of novel sequences, including potentially expressed novel genes and many previously unreported variants [6].

Another study identified 116 Mb of novel sequences from African cattle and illustrated how pangenomes can improve the precision of reference-guided alignment and variant detection compared to the current ARS-UCD1.2 reference assembly [7].

More recent pangenomic studies have further investigated NRUIs. A pangenome developed from 5 *de novo* genomes representing Indian dairy *Bos Indicus* breeds (including Gir, Kankrej, Tharparkar, Sahiwal, and Red Sindhi), identified 17.7 Mb of NRUIs [8]. The second study developed a pangenome comprising 64 *de novo* assemblies from 14 French cattle breeds, revealing 25 Mb of NRUIs [5]. Finally, the third study conducted a pangenome study in Holstein and Jersey breeds and identified, respectively, 63.75 and 42.34[Mb of novel sequences [9].

Together, these findings indicate that the organization of the cattle genome is more complex than previously believed and highlight the value of pangenome approaches to reveal functionally relevant SVs, particularly NRUIs, which contribute substantially to genomic diversity and the genetic architecture of key traits in cattle.

In the present study, we have constructed a cattle pangenome graph by integrating sixteen *de novo* haplotype-resolved genome assemblies originating from six major French dairy and beef breeds (Aubrac, Blonde d’Aquitaine, Charolais, Holstein, Montbéliarde, and Normande), three regional French dairy breeds (Abondance, Tarentaise, and Vosgienne) and yak (*B. grunniens*). Using this pangenome, we constructed a comprehensive catalog of SVs and NRUIs, performed functional annotation, and explored their potential contributions to economically important traits in cattle.

## Methods

### Sample collection

In this study, we constructed 14 haplotype-resolved genome assemblies using short and long read sequences derived from five pure bred [Abondance (ABO), Aubrac (AUB), Blonde d’Aquitaine (BAQ), Tarentaise (TAR) and Vosgienne (VOS)] and two cross-bred or hybrid trios [Holstein x Normande (HN) and Yak x Montbéliarde (YM)]. To enable a comprehensive bovine pangenome analysis, we also included the current ARS-UCD1.2 Hereford bovine reference genome [10] and two publicly available haplotype-resolved genome assemblies from the Charolaise breed [11].

### DNA extraction and sequencing

Genomic DNA was extracted from blood, semen, or ear biopsies following the protocol “DNA isolation from cattle tissues: blood, semen or any kind of tissues, including ear biopsies” [12] using Puregene kits (Qiagen) [13]. DNA quantity and quality were assessed using a Nanodrop One spectrophotometer (ThermoFisher Scientific).

For each parent of the trios, DNA libraries were prepared with the TruSeq Nano DNA HT Sample Preparation Kit (Illumina, CA, USA) in accordance with the manufacturer’s guidelines and then sequenced on an Illumina NovaSeq 6000 platform (Illumina, CA, USA) to produce 150-bp paired-end (PE) reads.

For the offspring, PacBio HiFi library preparation and sequencing were performed at the INRAE GeT-PlaGe core facility (Toulouse, France; https://get.genotoul.fr/la-plateforme/get-plage/) according to the manufacturer’s instructions “Procedure & Checklist Preparing HiFi SMRTbell Libraries using SMRTbell Express Template Prep Kit 2.0” [14].

### Construction of haplotype-resolved genome assemblies

Haplotype-resolved assemblies were generated using a trio binning approach, which partitions offspring long reads into haplotype-specific sets based on parental short reads [15]. Illumina paired-end (PE) short reads of the parents were used to create a kmer dictionary employing yak version 0.1 with the default parameters [16]. The offspring PacBio HiFi long reads were then assembled using Hifiasm version 0.19.8 [17] along with the parental kmer dictionaries (parameters −1 and −2) assign reads to their respective haplotypes. The resulting haplotype assemblies were scaffolded to near-chromosome level using RagTag scaffold tool version v2.1.0 with default parameters [18] aligned to the ARS-UCD1.2 Hereford reference (https://www.ncbi.nlm.nih.gov/datasets/genome/GCF_002263795.1/).

### Validation of parental haplotype-resolved assembly

The quality of the haplotype-resolved genome assemblies was evaluated using three metrics: total assembly length, N50 score, and completeness assessment. Assembly completeness of the assemblies was assessed using compleasm [19], which uses the miniprot protein-to-genome aligner in combination with conserved orthologous genes from BUSCO [20]. The assembly quality value (QV) was also quantified using Merqury version 1.3 [21]. In parallel, haplotypes were also aligned to the ARS-UCD1.2 reference genome using D-Genies 1.5 [22] with default parameters to visually inspect the dotplots for large-scale structural accuracy.

### Construction of a high-quality cattle pangenome graph

All 17 assemblies, including the ARS-UCD1.2 Hereford reference genome, were used to build a cattle pangenome graph with Minigraph (version 0-21) [23].

First, genetic distances between each of the 16 assemblies and the reference genome were estimated using Mash [24]. The pangenome graph was then built progressively, using the ARS-UCD1.2 reference as the backbone and incorporating the remaining assemblies in order of increasing genetic distance [see Additional file 2, Table S1]. To simplify the pangenome graph structure, only autosomes included in the construction.

### SV calling

The procedure for SV calling from the pangenome graph was conducted following the workflow described by Sorin *et al.* [5]. Briefly, allele information for each sample present in every bubble was retrieved by realigning each of the 16 assemblies back to the graph using the “-cxasm –call” option of Minigraph. Individual allele data were then merged into a single VCF file, following the guidelines set forth in the Minigraph-cookbook-v1 (https://github.com/lh3/minigraph) [23]. The resulting VCF file contains the final panel of SVs and records the presence or absence status of SVs in every sample.

### Recovery of Non-Reference Unique Insertion sequences (NRUIs)

NRUIs were extracted as outlined by Sorin *et al.* [5]. Briefly, three filters were applied: (1) nodes not present in any individual assembly paths were categorized as nested nodes and subsequently excluded from further analysis, (2) all nodes found in the reference path were also excluded, and (3) nodes corresponding to true bi-allelic insertions (defined as having a reference allele of zero length and an alternative allele exceeding 50 nucleotides) were retained as NRUIs.

### Population structure analysis using NRUIs

In order to capture a wider range of variability present in the *Bos taurus* genome, we expanded our NRUIs analysis to incorporate an additional panel of NRUIs derived from a previous a pangenome built with 64 *de novo* assemblies [5]. The final panel of NRUIs consisted of 45,136 NRUIs covering a total length of 48,202,473 nucleotides.

To assess NRUI distribution across a larger French bovine population, we used the Minigraph tool to realign a total of 181 *de novo* assemblies from 14 French cattle breeds and yak (comprising 165 publicly available *de novo* assemblies retrieved from projects PRJEB64023 and PRJEB59364, along with the 16 haplotype-resolved assemblies of this study). These assemblies were aligned onto both the pangenomes we developed in this study and the one previously created with 64 *de novo* assemblies [5].

From the pangenome alignments, a pathway was established for each assembly and the genotype of each animal was predicted for every NRUI in the merged panel. Genotypes for NRUIs common to both panels were extracted from the alignments on the current pangenome, while genotypes or NRUIs specific to the previous panel were obtained from the alignments on the previous pangenome. Subsequently, both sets of genotypes were merged into a single VCF file. A binary presence/absence matrix was generated for each animal and used for hierarchical clustering analysis using the hclust function of the R package [25].

### Functional characterization of NRUIs

To investigate the potential functional impact of NRUIs on cattle phenotypes, two complementary analyses were performed.

First, cattle quantitative trait loci (QTL) data from the publicly accessible Animal QTL database [26] with NRUI genomic regions and enrichment analyses were conducted using the GALLO R package [27]. Traits with an FDR-adjusted p-value < 0.001 were considered significantly enriched.

In the second study, the cattle gene annotation GTF file (Ensembl release 110; https://ftp.ensembl.org/pub/release-110/gtf/bos_taurus/Bos_taurus.ARS-UCD1.2.110.gtf.gz) was retrieved. NRUIs overlapping exons, coding sequences (CDS), introns, and untranslated regions (UTRs) were identified, and the corresponding genes were subjected to gene ontology (GO) enrichment analyses using the PANTHER Classification System [28].

### Effect of selected NRUIs on recorded traits

The potential phenotypic impact of NRUIs was evaluated using a sequential approach. First, the SV catalogue was filtered to identify NRUIs that were breed-specific and located within annotated gene regions, including both coding and non-coding sequences.

These NRUIs were then further refined using functional annotations, that were obtained in the previous step, to retain only those within known genes associated with milk production and developmental processes. From this filtered set, two candidate NRUIs, specific to the Normande breed and located within the intronic regions of *ARMH3* and *EPHA5*, were selected for proof-of-concept analyses.

To investigate the impact of these two NRUIs on key cattle phenotypes, we applied a haplotype-based testing strategy using a large cohort of animals with available phenotypes and genotypes derived from various low- or medium-density SNP arrays. Long-read sequences for 28 Normande samples were retrieved from the European Nucleotide Archive (projectID PRJEB68295 and PRJEB59364), and realigned to the ARS-UCD1.2 reference genome using the PacBio pbmm2 aligner [29]. SVs were then called using the PacBio pbSV SV caller [30], allowing us to determine the true genotypes of the 28 samples for the two NRUIs of interest [see Additional file 2, Table S2].

After identifying NRUI carriers (cases) and non-carriers (controls) based on sequence-derived genotypes, we screened a panel of 241,413 Normande animals genotyped for routine genomic evaluation to identify haplotypes of markers that exhibit strong linkage disequilibrium (LD) with the candidate NRUIs. Two filters were applied: (1) all cases had to possess the haplotype, and (2) the number of controls having this haplotype was minimized. These two filters substantially decreased the number of false positives (*i.e.*, individuals carrying the haplotype but not the NRUI). Although a limited number of false negatives persisted (NRUI carriers classified as controls), the LD between the haplotypes and the candidate NRUIs was high, ensuring reliable subsequent analysis.

In order to evaluate the haplotype-based prediction strategy and to confirm the concordance between the haplotype status and true genotypes, the NRUI insertion positions were confirmed through PCR amplification and agarose gel electrophoresis. DNA from six animals were used to performed this cross-validation: two heterozygotes, two homozygous carriers, and two non-carrier. Primers were designed using primer3 version 4.1.0 [31–33] (Table 1) and synthesized by Eurofins Genomics (Ebersberg, Germany). They were specifically designed to span the NRUI breakpoints, producing two amplicons of different sizes corresponding to each allele. PCR amplification was performed using previously described protocol [5] and PCR products were analyzed through gel electrophoresis on a 2% agarose gel. For each NRUI, non-carriers were expected to exhibit the shortest amplicon, homozygous carriers the largest amplicon, and heterozygotes both types of amplicons.

**Table 1.**
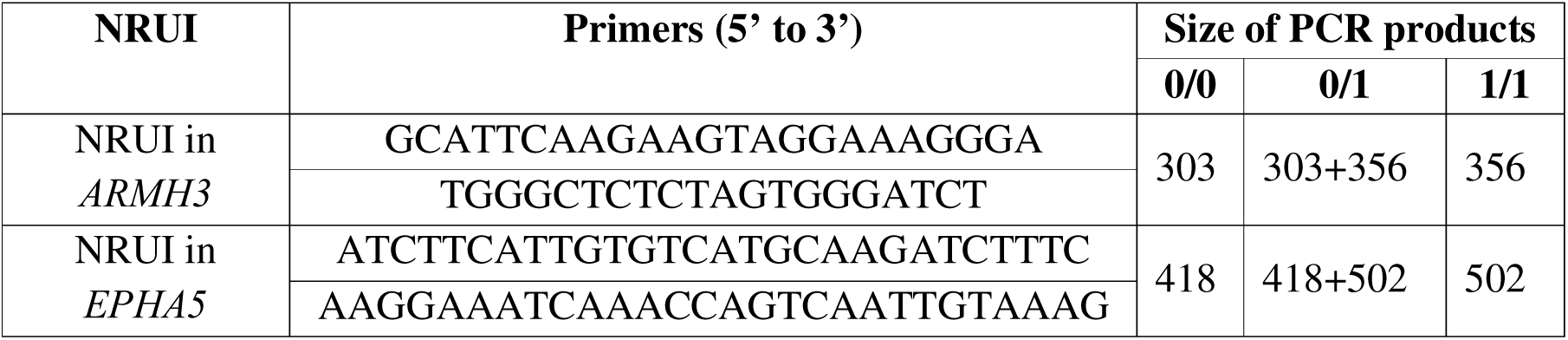
Features of PCR primers.

Sanger sequencing was also performed on DNA from one animal homozygous for the haplotype. The PCR product was sequenced by Eurofins Genomics (Ebersberg, Germany) to determine the exact genomic insertion location of each examined NRUI.

The effects of the haplotypes in heterozygous and homozygous states were estimated for 14 morphological and production traits routinely recorded for selection purposes as described in Besnard *et al.* [34]. To account for environmental factors influencing these phenotypes, we used yield deviation (YD) data, *i.e.*, phenotypes adjusted for non-genetic effects as estimated in the national genomic evaluation models. YD are a by-product of the official genomic evaluations carried out by GenEval for French breeding organizations [35].

YD were analyzed using the blupf90 family of programs, specifically renumf90 followed by blupf90. A mixed model was fitted with haplotype and year of birth as fixed effects, and an animal random polygenic effect. Overall, sample sizes ranged from 13,990 cows for teat length and rear teat placement to 119,842 cows for heifer non-return rate [see Additional file 2, Table S3].

Trait means between group of genotypes were compared using Student’s t-test, and p-values were adjusted for multiple testing using the Benjamini-Hochberg correction in R software. To facilitate comparisons across traits, haplotype effects were expressed in genetic standard deviations (GSD), as estimated for the national genomic evaluations.

### Prediction of novel functional features in NRUIs

To identify novel functional elements in NRUI sequences, yak-specific NRUIs were excluded and then repetitive sequences were masked using RepeatMasker v4.1.5 with the parameter option “-species cow” [36]. *Ab initio* gene structure prediction was then carried out on the masked sequences using the Augustus version 3.3.3 tool [37] with default parameters trained on the human genome. This facilitated the identification of NRUIs that contained potential gene structures. Amino acid sequences of the predicted genes were extracted using the “getAnnoFasta.pl” perl script from the Augustus package and subsequently compared against the cattle protein database to identify reference-liked gene coding proteins.

In parallel, masked NRUI sequences were compared against a multi-species protein database comprising three Bovidae (*Bos taurus, Bos grunniens, Bos mutus*), 11 other mammalian species (*Capra hircus, Ovis aries, Bison bison bison, Camelus dromedaries, Equus caballus, Delphinapterus leucas, Balaenoptera musculus, Homo sapiens, Mus musculus, Physeter catodon, and Tursiops truncatus*), along with the curated protein databases of SwissProt (https://ftp.ncbi.nlm.nih.gov/blast/db/FASTA/swissprot.gz) and PDB (https://ftp.ncbi.nlm.nih.gov/blast/db/FASTA/pdbaa.gz). Comparisons were performed using BlastX implemented in the Diamond-version 0.9.30 tool [38] enabling the identification of both reference-liked protein-coding genes and novel genes encoding proteins known in other species but not yet annotated in cattle.

Finally, the expression profiles of the predicted novel genes were assessed using a total of 71 raw RNA-seq data from blood and milk cells downloaded from the NCBI Short-Read Archive database [see Additional file 2, Table S4]. This analysis enabled the validation of these predicted genes as truly expressed genes in the species under study. The raw RNAseq data were processed with the nf-core/RNAseq pipeline (https://nf-co.re/rnaseq/3.21.0/), aligned to the NRUI sequences using STAR [39] and quantified using RSEM [40] along with the GTF gene annotation output file produced by Augustus.

## Results

### Construction of haplotype-resolved genome assemblies

Sequencing of the seven cattle samples yielded a total of 793 Gb of raw PacBio HiFi long-read sequences, achieving an average sequence depth of 41X per genome. For each sample, two haplotype-resolved genome assemblies were constructed; resulting in the production of 14 assemblies with an average scaffold N50 value of 89.9 Mb and an average L50 value of 13 scaffolds (Table 2). The mean QV score was 61.5 and ranged from 54.5 for the VOS_hap1 up to 70.3 for Yak_hap1 (Table 2). Also, the mean assembled genome size was 3.03 Gb and ranged from 2.7 Gb for the ABO_hap1 assembly to 3.26 GB for the VOS_hap2 assembly. Together, these metrics indicate that the genome assemblies were highly continuous and displayed minimal fragmentation. The mean completeness of genome assemblies was 95.4%, with individual completeness scores varying from 86.7% to 98.8%, confirming their high overall quality. Finally, the alignment of these assemblies with D-Genies demonstrated a high degree of similarity and concordance with the chromosomes of the ARS-UCD1.2 reference genome assembly [see Additional file 1, Fig. S1-S8].

**Table 2:**
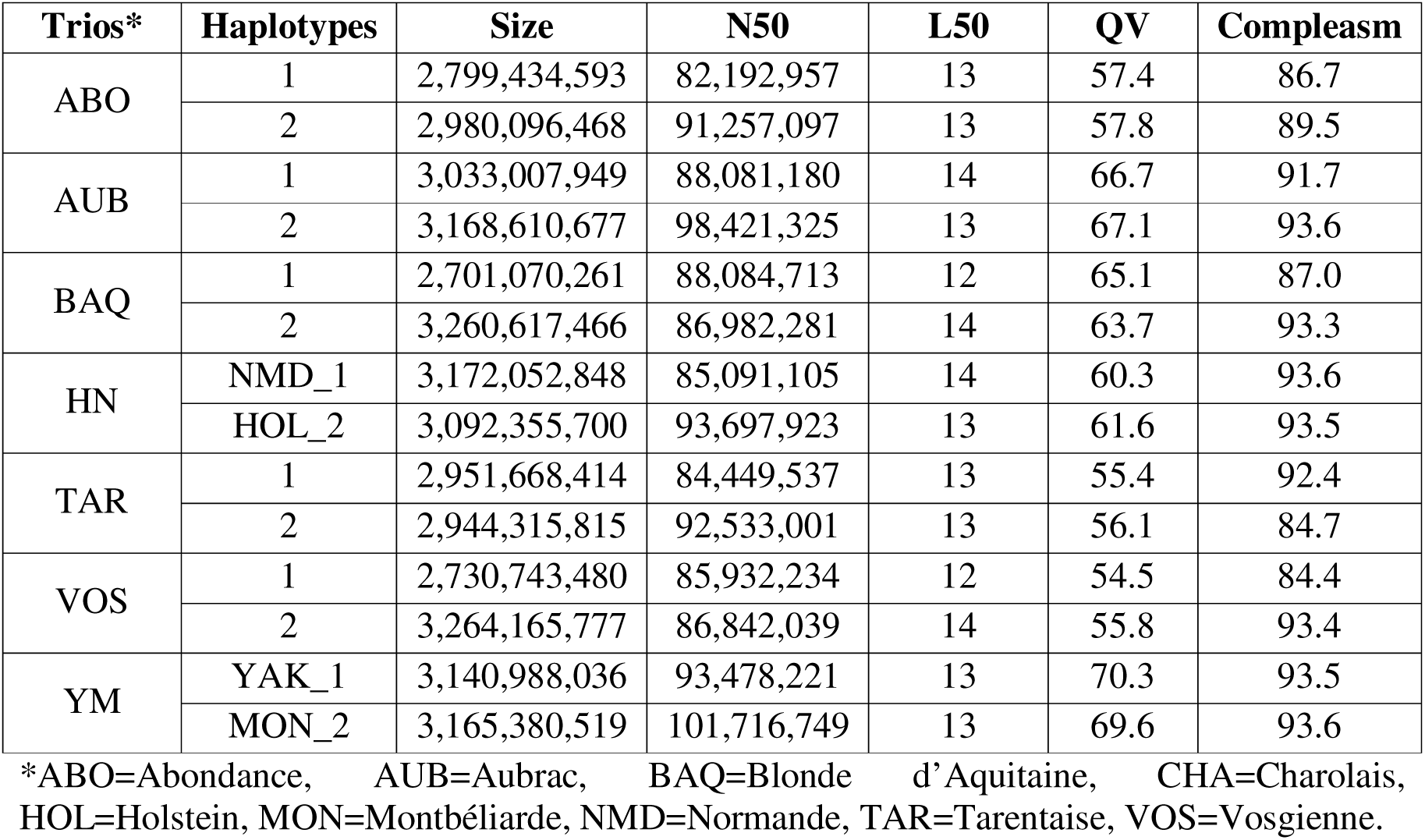
Metrics of the 16 assemblies used to construct the pangenome graph.

| **Trios\*** | **Haplotypes** | **Size** | **N50** | **L50** | **QV** | **Compleasm** |
| --- | --- | --- | --- | --- | --- | --- |
| ABO | 1 | 2,799,434,593 | 82,192,957 | 13 | 57.4 | 86.7 |
|  | 2 | 2,980,096,468 | 91,257,097 | 13 | 57.8 | 89.5 |
| AUB | 1 | 3,033,007,949 | 88,081,180 | 14 | 66.7 | 91.7 |
|  | 2 | 3,168,610,677 | 98,421,325 | 13 | 67.1 | 93.6 |
| BAQ | 1 | 2,701,070,261 | 88,084,713 | 12 | 65.1 | 87.0 |
|  | 2 | 3,260,617,466 | 86,982,281 | 14 | 63.7 | 93.3 |
| HN | NMD_1 | 3,172,052,848 | 85,091,105 | 14 | 60.3 | 93.6 |
|  | HOL_2 | 3,092,355,700 | 93,697,923 | 13 | 61.6 | 93.5 |
| TAR | 1 | 2,951,668,414 | 84,449,537 | 13 | 55.4 | 92.4 |
|  | 2 | 2,944,315,815 | 92,533,001 | 13 | 56.1 | 84.7 |
| VOS | 1 | 2,730,743,480 | 85,932,234 | 12 | 54.5 | 84.4 |
|  | 2 | 3,264,165,777 | 86,842,039 | 14 | 55.8 | 93.4 |
| YM | YAK_1 | 3,140,988,036 | 93,478,221 | 13 | 70.3 | 93.5 |
|  | MON_2 | 3,165,380,519 | 101,716,749 | 13 | 69.6 | 93.6 |

\*ABO=Abundance, AUB=Aubrac, BAQ=Blonde d'Aquitaine, CHA=Charolais, HOL=Holstein, MON=Montbéliarde, NMD=Normande, TAR=Tarentaise, VOS=Vosgienne.

### Construction of a bovine pangenome graph

Integration of the 16 haplotype-resolved assemblies using the autosomal sequences of the Hereford ARS-UCD1.2 reference genome as a backbone produced a bovine pangenome graph with 273,192 nodes linked by 388,858 edges, totalling 2,569,341,538 nucleotides. The core component of the pangenome consisted of 101,248 nodes spanning 2,435,570,613 bases, whereas the flexible region comprised 170,648 nodes corresponding to 132,903,963 nucleotides. A small subset of 1,296 nodes, representing 866,962 nucleotides, could not be traversed by any of the assemblies. These likely reflect limitations in accurately mapping certain complex graph structures rather than true biological absence. Consistent with previous reports [5]. Such unresolved nodes primarily occurred within highly nested or structurally complex bubbles in the graph and were therefore excluded from downstream analyses.

### Discovery of SVs from the pangenome graph

Analysis of the pangenome graph identified 101,219 SVs, of which 95,137 variants (94%) were classified as biallelic, while only 6,082 SVs (6%) were found to be multi-allelic. Biallelic SVs were predominantly composed of insertions (45.3%) and deletions (35.1%), while the remaining SVs (19.6%) corresponded to sequence substitutions, defined by the presence of non-zero sequence lengths for both reference and alternate allele (Table 3).

**Table 3:** Distribution of SVs by allele count and SV type.

| Mutations | Bi-allelic SVs |  | Multi-allelic SVs | Total |
| --- | --- | --- | --- | --- |
| Insertions | 43,096 | 33,855* | 609 | 43,705 |
|  |  | 9,241** |  |  |
| Deletions | 33,422 | 26,286* | 778 | 34,200 |
|  |  | 7,136** |  |  |
| Others variants *** | 18,619 |  | 4,695 | 23,314 |
| **Total** | **95,137** |  | **6,082** | **101,219** |

\*Alternative allele sequence length = 0 for deletion and reference allele sequence length = 0 for insertions; \*\* alternative allele sequence length between 1 and 5 nucleotides for deletion and reference allele sequence length between 1 and 5 nucleotides for insertions; \*\*\* all others SVs not classified as insertions nor deletions.

To further characterize the SVs, we analysed the size distribution of insertions and deletions and observed very similar profiles. The vast majority of variants were small, ranging from 50 to 2,000 nucleotides (Fig. 1). Furthermore, both deletions and insertions exhibited distinct peaks at approximately 150-300 bp, 1.5 kb, and 8.5 kb, which are likely due to the activity of specific transposable element classes of SINES (mainly Bov_A2 at the 150-300 bp peak) and LINEs (mainly BovB at the 1.5 kb peak; and the L1_BT class at the 8.5kb peak).

**Figure 1.**
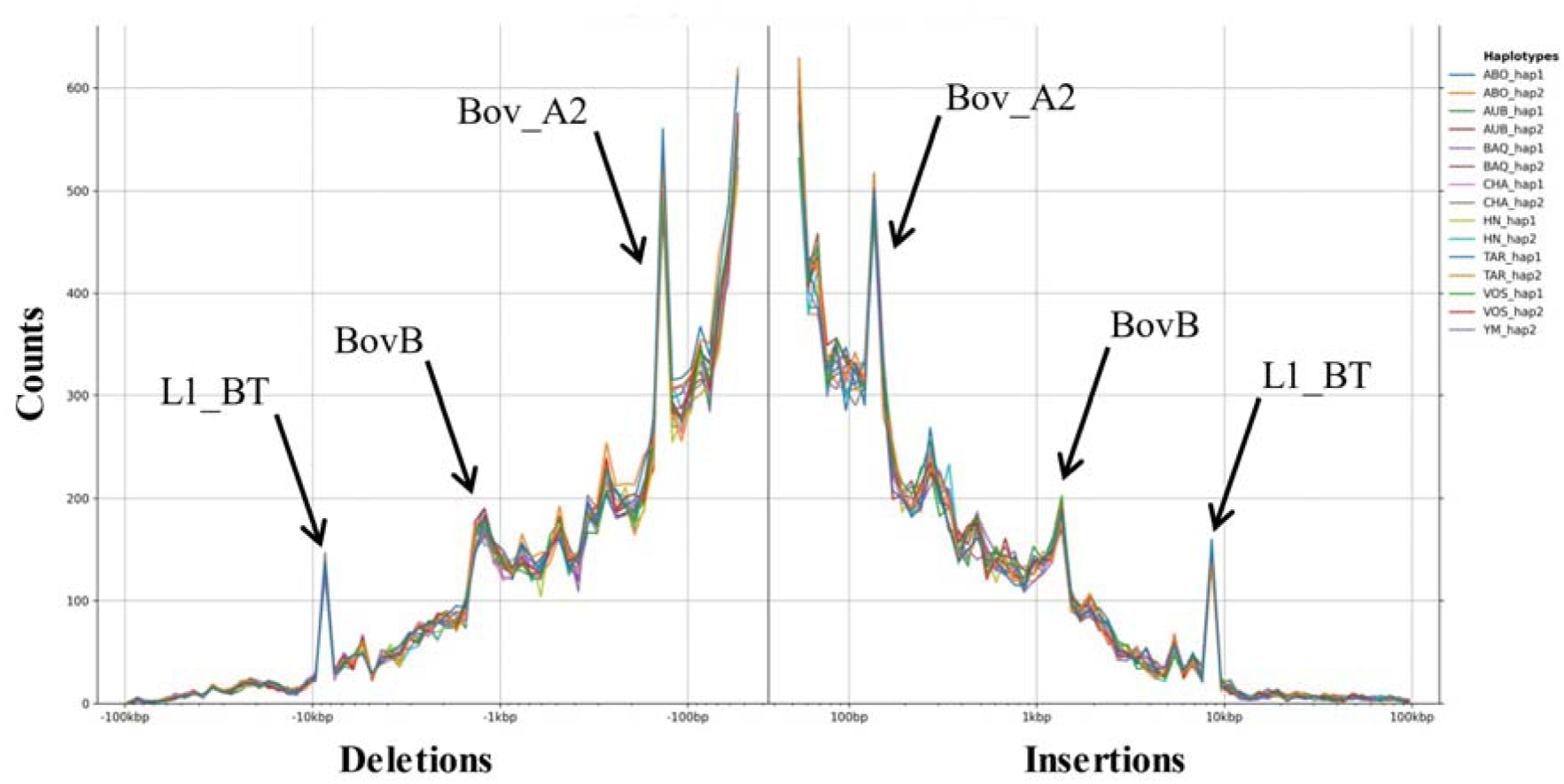
Size distribution of INS and DEL Size distribution of SVs classified as deletions (left) and insertions (right), identified using Minigraph across the 16 phased haplotypes. The arrows highlight peaks at 150-300bp, 1.5 kb and 8.5 kb, which are likely driven by the activity of specific transposable element classes

### Recovery of NRUIs from the pangenome graph

Examination of true insertions from biallelic bubbles, excluding reference genome nodes, identified 33,634 non-reference unique insertions (NRUIs), totalling 34,405,480 bases. Of these, 16,614 NRUIs were specific to yak, spanning 14,663,556 bases, while the remaining 17,020 NRUIs were present in at least one *Bos taurus* breed, covering 19,741,924 bases.

Comparison of the 17,020 *Bos taurus* NRUIs with the panel constructed in our previous pangenome study based on 64 *de novo* genome assemblies revealed the identification of 9,481 common unique insertions (55.6%) representing 6,038,185 nucleotides (30.6%). Out of these, 94.4% (8,950 NRUIs covering 5,542,808 bases) showed identical insertion coordinates in both panels.

Because these two panels were developed from a genetically diverse populations of French cattle, we merged these NRUIs to form a single panel, allowing us to capture a wider spectrum of genetic diversity found within the genome of French cattle breeds. The resulting merged NRUI panel contained 28,998 unique insertions spanning a total length of 33,742,494 nucleotides. This panel was subsequently used to genotype a broader population consisting of 181 animals originating from 14 French cattle breeds and then investigate their frequencies at the population level.

The vast majority of NRUIs had a low frequency within the studied population (Fig. 2A). Approximately 42% had frequencies of 5% or less. Conversely, 124 NRUIs have been observed in at least 99% of the animals but were absent from the reference genome. These likely reflects deletions specific to the Hereford genome or to the animal (Dominette) used to construct the reference genome sequence.

**Figure 2.**
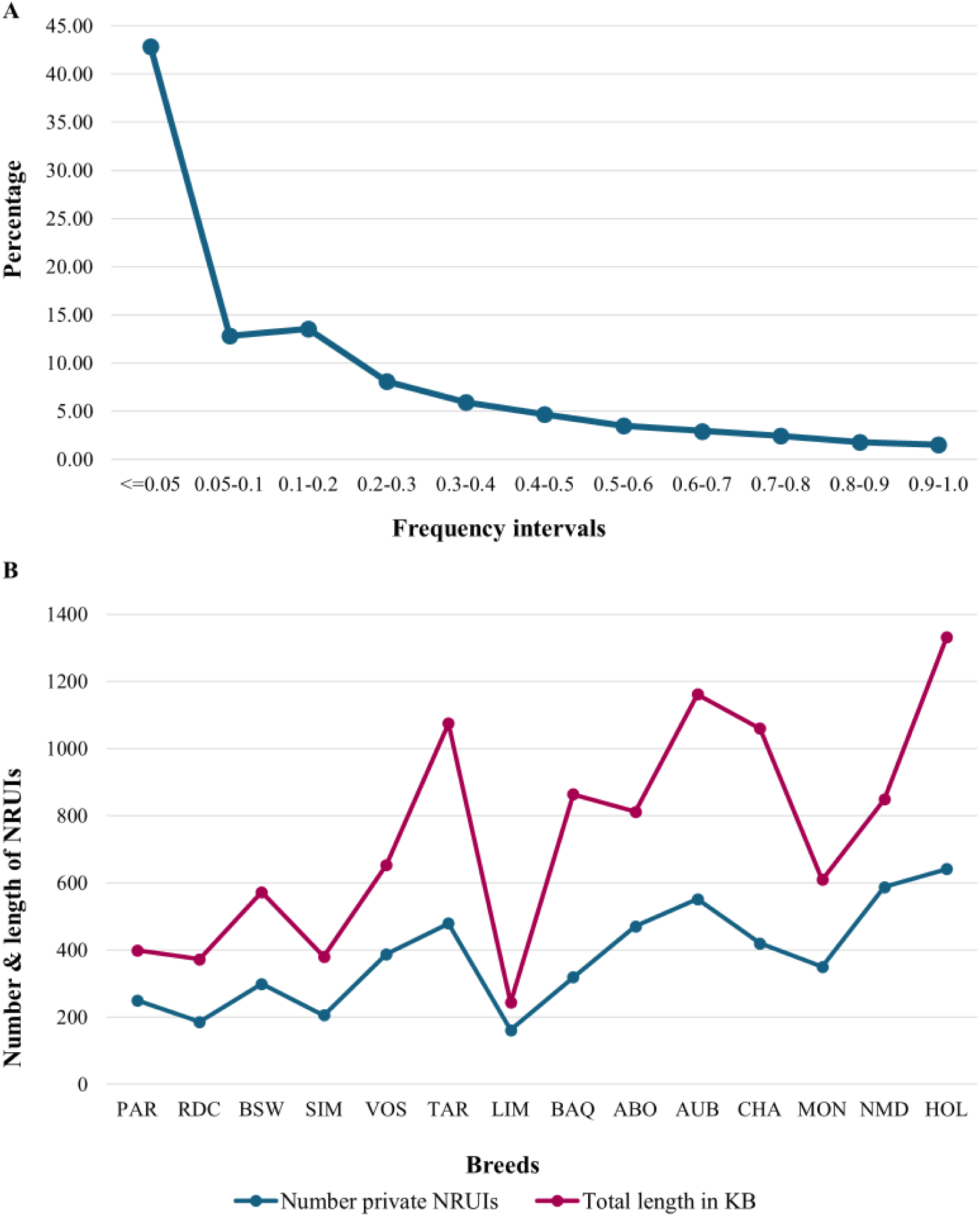
Distribution of NRUIs within the cattle breeds (A) Distribution of NRUI frequencies at the population level. (B) Distribution of the private NRUIs. *ABO=Abondance, AUB=Aubrac, BAQ=Blonde d’Aquitaine, CHA=Charolais, HOL=Holstein, MON=Montbéliarde, NMD=Normande, TAR=Tarentaise, VOS=Vosgienne.

Furthermore, 5,311 NRUIs were found to be private as they were observed in only one breed (Fig. 2B). These private NRUIs covered a total of 10.4 Mb, with breed-specific totals ranging from 243.8 kb in Limousine to 1.4 Mb in Holstein.

### Functional characterization of NRUIs

The merged panel of NRUIs was subsequently used to characterize their functional properties by performing Gene Ontology (GO) enrichment analysis for genes whose exonic, intronic, or regulatory regions overlapped with NRUIs, and QTL enrichment and association analyses to explore their impact on economically important cattle traits.

Most NRUIs (69.1%) were identified in intergenic regions. The remaining 8,951 NRUIs (30.9%) were located within 4,348 unique cattle gene regions. Among these, 8,641 NRUIs (29.8%) were found within intronic regions, while only 212 NRUIs (0.73%) were identified within exons of 201 genes and 98 NRUIs (0.34%) were located within the 5’ or 3’ UTR regions of 91 unique genes.

Gene enrichment analyses using Panther classification [28] revealed that the many genes affected by NRUIs were unclassified as no Panther category was assigned. Among the classified genes, several GO molecular function terms [see Additional file 2, Table S5] were found to be significantly over-represented, namely *binding* (GO:0005488), *catalytic activity* (GO:0003824), *transporter activity* (GO:0005215), molecular function regulator activity (GO:0098772), and transcription regulator activity (GO:0140110) (Fig. 3A). The three most significant protein classification categories corresponded to metabolite interconversion enzymes (PC00262), protein modifying enzyme (PC00260), and transporter class (PC00227) (Fig. 3B) [see Additional file 2, Table S6]. Finally, pathway analysis identified Wnt signalling pathway (P00057), and Gonadotropin-releasing hormone receptor pathway (P06664) as the two most over-represented pathways (Fig. 3C) [see Additional file 2, Table S7].

**Figure 3.**
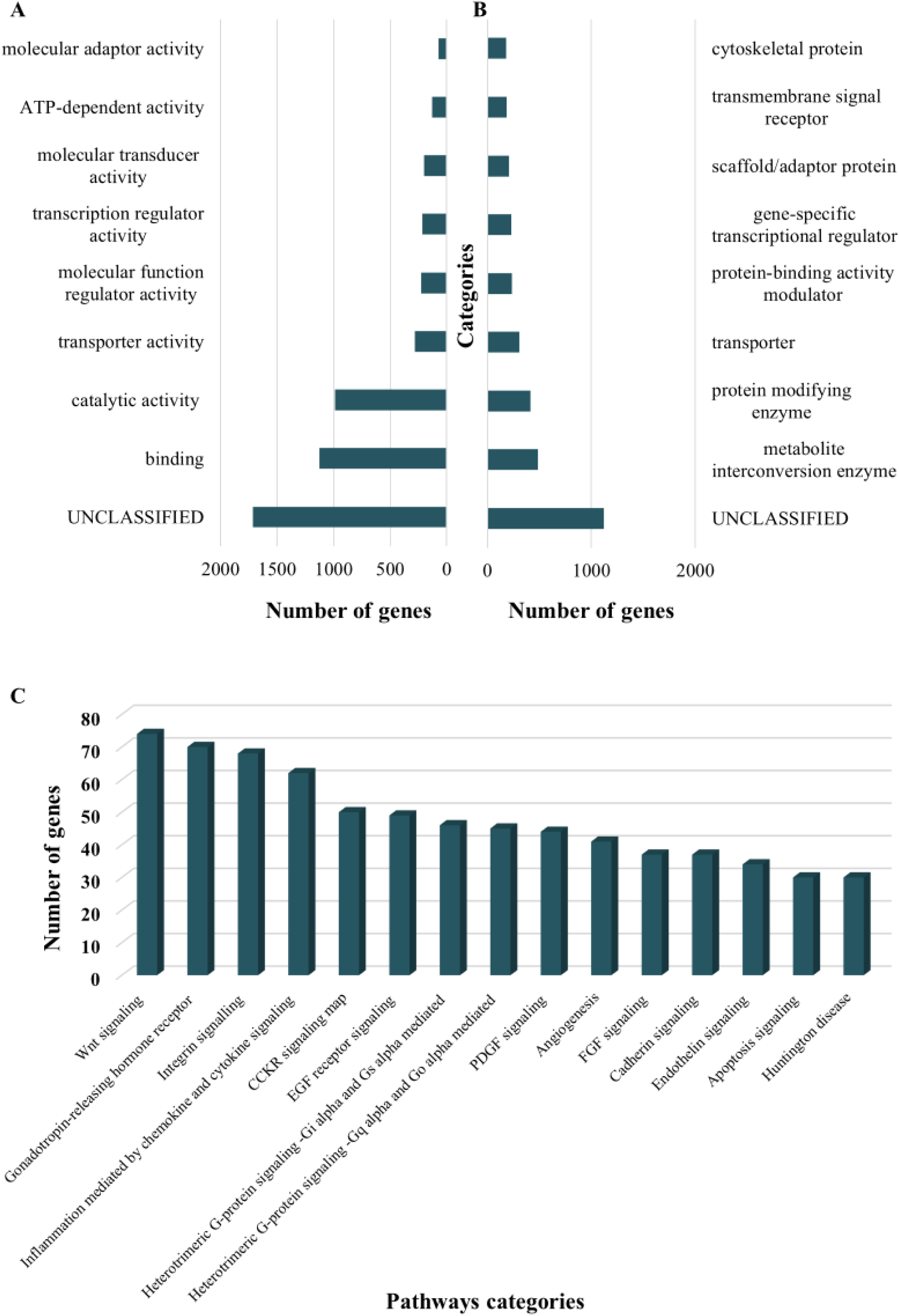
Functional implications of SVs (A) Molecular enrichment analysis of the NRUIs panel. (B) Protein classes enriched in the NRUIs panel. (C) Representation of the 15 most enriched pathways in the NRUIs panel.

QTL enrichment analyses revealed that NRUIs were significantly enriched in QTL regions associated with 24 economically important traits in cattle (Fig 4). The majority of NRUIs overlapping QTL were related to milk production (59.5%), with the most significant associations observed for milk fat percentage (26.0%), followed by milk protein percentage (21.3%), and milk yield (16.3%).

**Figure 4.**
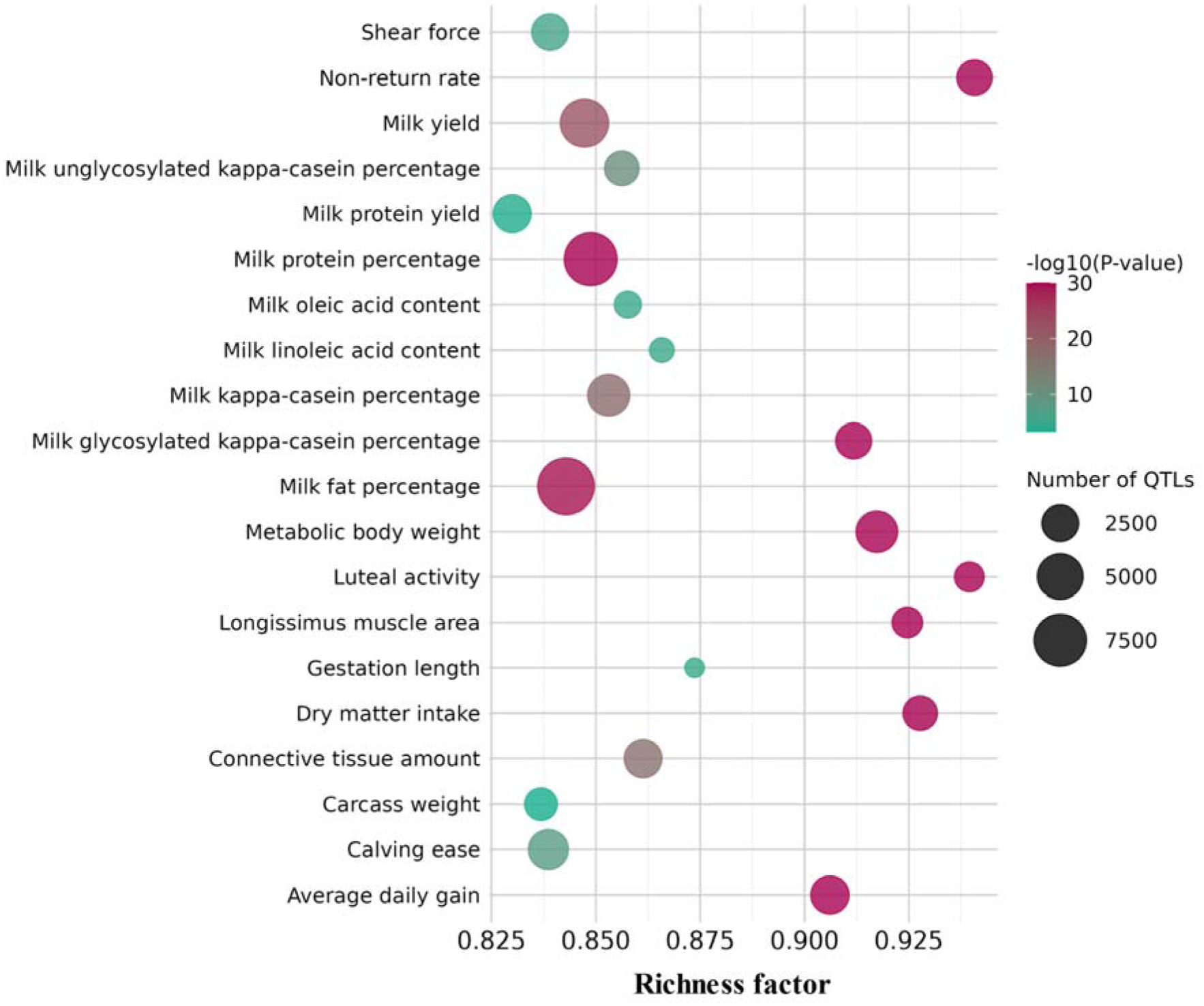
QTL enrichment analysis of NRUIs Bubble plot of the enrichment analysis of the top 20 QTLs impacted by NRUIs.

### Impact of NRUIs on cattle key phenotypes

Combined gene annotation efforts along with QTL enrichment and analyses of the distribution of allele frequencies within breeds enabled us to identify 1,660 breed-specific NRUIs located within candidate genes. Of particular interest, two NRUIs have been found to be specific yet polymorphic (MAF=9%) within the Normande breed, and were located within the intronic regions of two genes of interest, namely *ARMH3* and *EPHA5*. The first NRUI is 53 nucleotides in length and was inserted within intron 22 (out of 25 introns) of the *ARMH3* gene. The second NRUI is 84 nucleotides in length and was located within intron 3 (out of 16 introns) of the *EPHA5* gene. The exact genomic positions of these two NRUIs were further validated by Sanger sequencing of the flanking regions: the *ARMH3* NRUI was located on BTA26 at position 22,672,107 (Fig. 5B) and the *EPHA5* one on BTA6 at position 81,086,501 (Fig. 5DC).

**Figure 5.**
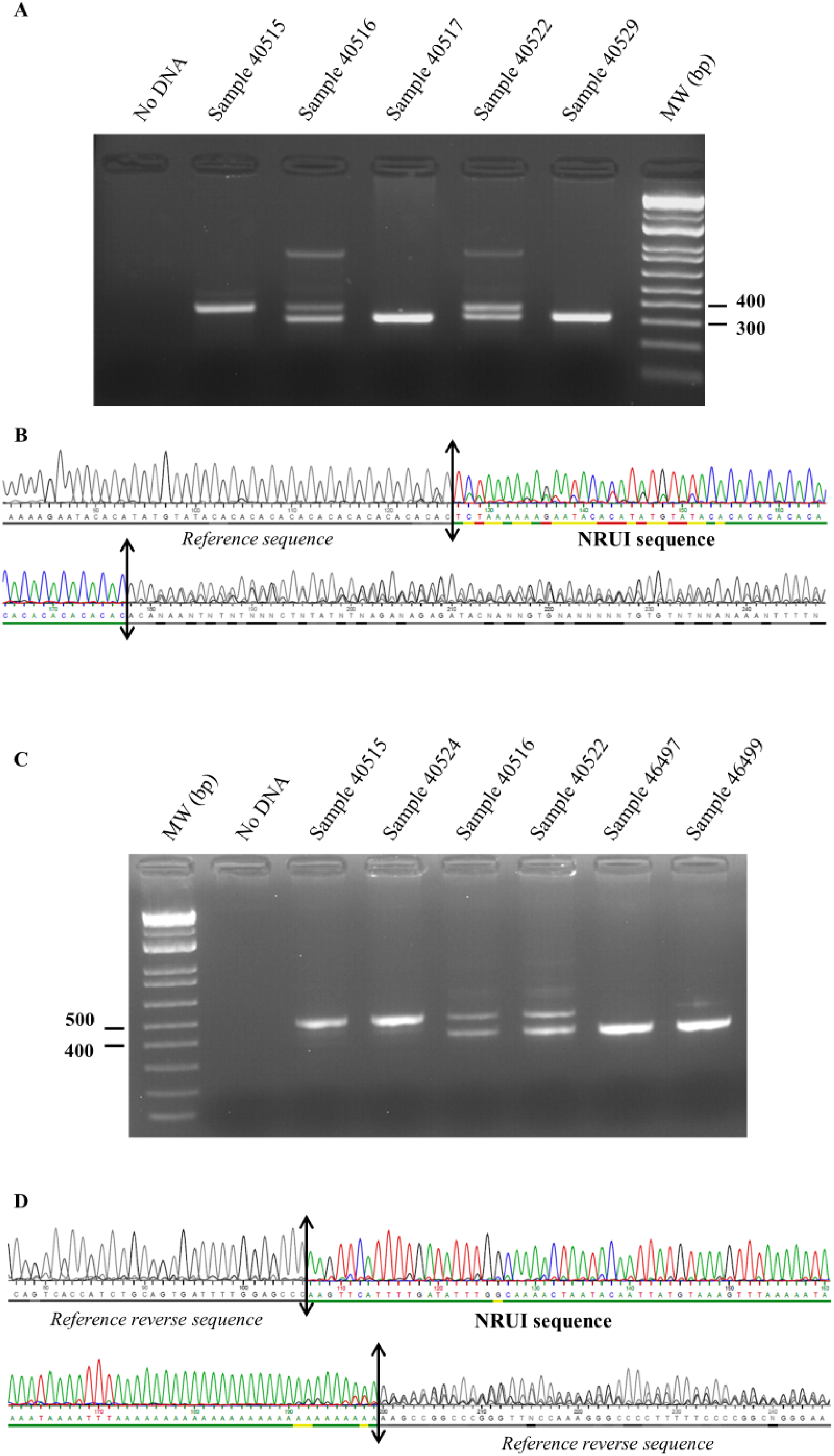
PCR and sequencing validation of the two Normande NRUIs (A) Gel electrophoresis of 5 samples for *ARMH3* NRUI: one homozygous reference sample (40515), two heterozygous samples (40516 and 40522), and two homozygous samples for the NRUI (40517 and 40529). (B) Sanger sequencing of the *ARMH3* NRUI sequence and its breakpoints. (C) Gel electrophoresis of 6 samples for *EPHA5* NRUI: two homozygous reference samples (40497 and 46499), two heterozygous samples (40516 and 40522), and two homozygous samples for the NRUI (40515 and 40524). (D) Sanger sequencing of the *EPHA5* NRUI sequence and its breakpoints.

Both genes have been previously linked to important cattle phenotypes. In this study, we explored their potential impact on several economic traits. To achieve this, we first identified the haplotypes that were in complete LD with the candidate NRUIs. To evaluate the quality of these haplotypes, we compared their status in six selected animals (2 homozygous reference, 2 heterozygous, and 2 homozygous alternate) against their true genotype, which was determined through PCR amplification of their DNA (Fig. 5A and C). The findings from this comparison indicate that the genotype of each animal, derived from its DNA, aligns perfectly with its status. This led to the conclusion that the haplotypes exhibit strong LD with the two variants, and that carriers of these haplotypes are also carriers of the NRUIs.

Following this, we performed a Student’s t-test to investigate potential associations between these two NRUIs and significant phenotypes in cattle. The NRUI in the intronic regions of *ARMH3*, showed a clear dosage-dependent pattern for stature, with heterozygotes increasing height by 0.25 GSD and homozygotes by 0.50 GSD (Fig. 6A). Additive effects were also detected for chest depth (0.18 GSD) and udder depth (0.25 GSD) (Fig. 6B), as well as milk protein (−0.17 GSD) and fat content (−0.12 GSD) (Fig. 6C and D). No significant effect on milk yield was observed [see Additional file 2, Table S3].

**Figure 6.**
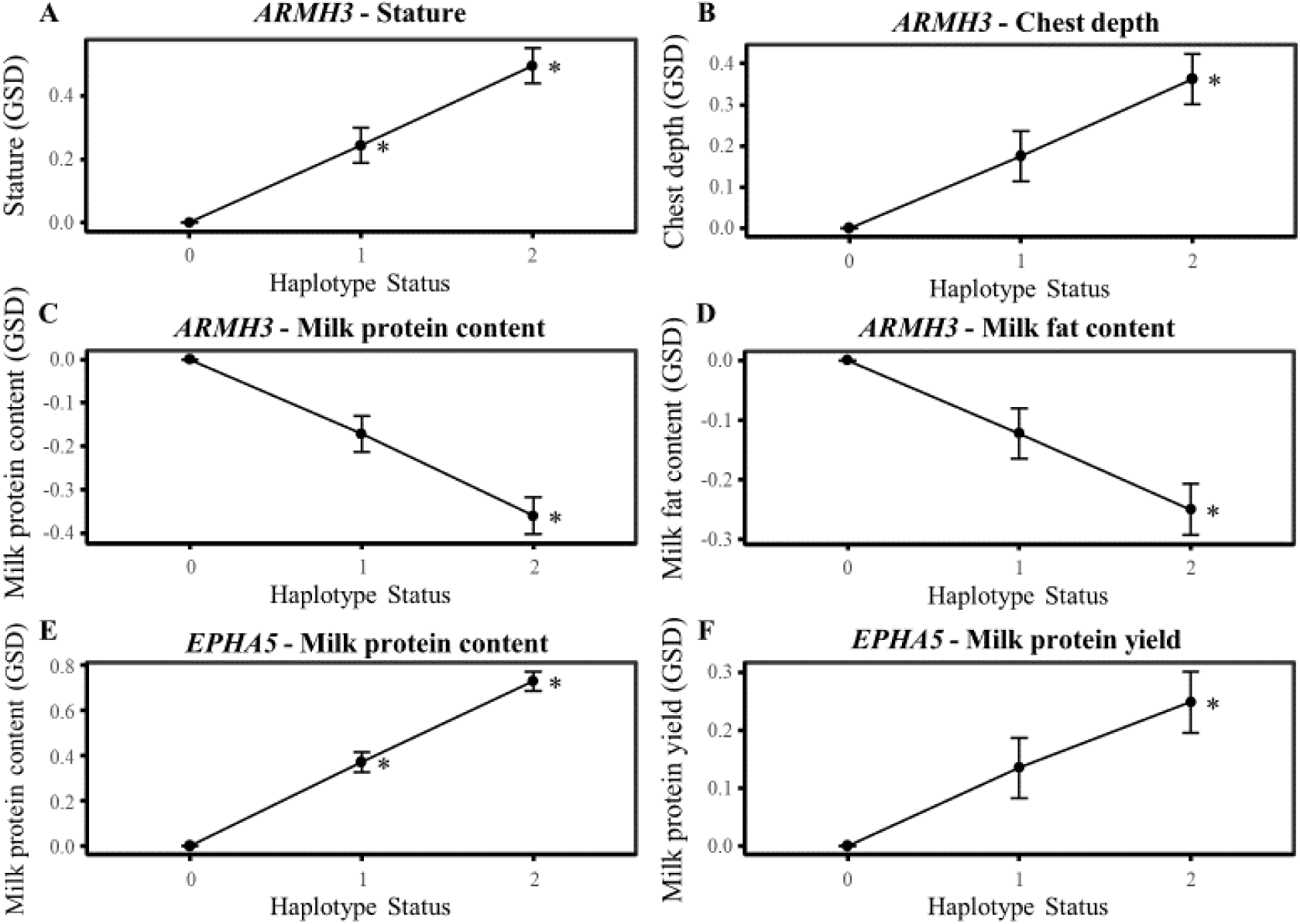
Impact of NRUIs on cattle phenotypes (A) Effect of *ARMH3* NRUI on stature trait. (B) Effect of *ARMH3* NRUI on chest depth trait. (C) Effect of *ARMH3* NRUI on milk protein content trait. (D) Effect of *ARMH3* NRUI on milk fat content trait. (E) Effect of *EPHA5* NRUI on milk protein content trait. (F) Effect of *EPHA5* NRUI on milk protein yield trait. *GSD*: genetic standard deviations, 0: homozygous for reference allele, 1: heterozygous, and 2: homozygous for the NRUI allele.

For the NRUI located with the *EPHA5* intronic regions, a strong additive effect was observed on milk composition traits. The effect on milk protein content was particularly pronounced, reaching 0.37 GSD at the homozygous state (Fig. 6E). We also observed a moderate but consistent additive effect on milk protein yield (0.13 GSD), which was only significant at the homozygous state (Fig. 6F). A smaller yet positive effect was also detected on milk fat content (−0.14 GSD at the homozygous state) [see Additional file 2, Table S3].

### Prediction of novel functional elements in NRUIs

Repeat annotation of NRUIs revealed that 45.34% (15.3 Mb) were repeated sequences [see Additional file 2, Table S8]. Most of these were interspersed repeats (39.9%), which primarily consist of long interspersed nuclear elements (LINEs, 36.5%), long terminal repeats (LTR, 2.25%), DNA elements (0.7%), and short interspersed nuclear elements (SINEs, 0.5%).

Functional annotation of NRUIs using Augustus gene structure prediction identified a total of 923 NRUIs that encompassed 970 novel genes, of which 429 were predicted to be complete gene models with an average size of 310 amino acids and ranging from to 67 up to 1,182 amino acids. For these complete gene models, a transcription start site, a start codon, at least one exon, one stop codon, and a transcription termination site were predicted. Among the 429 complete genes, 173 demonstrated homology to known bovine proteins (bovine-like genes), whereas 251 genes showed homology with proteins from at least one species other than *Bos taurus* (true novel genes). Five complete genes had no detectable homology [see Additional file 2, Table S11]. Among the five genes, two were identified as being expressed in at least one RNA-Seq sample, qualifying them as true novel genes. The first gene displayed a TPM of 19.91, while the second gene was expressed across five RNA-Seq samples, with a TPM varying from 5.43 to 11.73 [see Additional file 2, Table S11]. In contrast, the remaining three genes did not exhibit any expression profiles in the RNA-Seq samples. This lack of expression may indicate either false predictions by Augustus or that these genes are not expressed in the tissues from which the 71 RNA samples were derived.

Expression validation using 71 RNA-seq datasets demonstrated that 802 of the 970 genes (82.6%) were expressed in at least one of the RNA-Seq samples [see Additional file 2, Table S10]. Among these, 290 genes (∼36.2%) were expressed in over 90% of the samples, suggesting their potentially significant role in cellular processes, while 155 genes (19.3%) were expressed in fewer than 10% of the samples, indicating potential involvement in specific or rare biological processes.

## Discussion

In comparison to linear reference genome sequences, graph-based pangenomes facilitate a more comprehensive detection and representation of genomic sequences and variations, resulting in improved read alignment rates with the reference and enhancing the ability to capture missing heritability within populations.

In this study, we constructed a cattle pangenome graph using 16 haplotype-resolved genome assemblies originating from eight French cattle breeds (Abondance, Aubrac, Blonde d’Aquitaine, Charolais, Holstein, Montbéliarde, Normande, Tarentaise, and Vosgienne) and a closely related species, yak. We focused solely on autosomal sequences since the haplotype-resolved assemblies depict either paternal or maternal haplotypes, thereby excluding sex chromosomes (X and Y), mitochondrial DNA, as well as the unplaced contig chromosomal sequences.

The resulting bovine pangenome graph encompassed 2.57 Gb distributed across 273,192 nodes interconnected by 388,858 edges. The core genome represented 94.7% (2.44 Gb) of the graph whereas the flexible region accounted for only 5.1% (133 Mb) of the pangenome. These results were very similar to those previously reported by two independent pangenome studies that also considered only autosomes and used Minigraph to build the bovine pangenome [3][6]. However, our current pangenome exhibited a significantly lower number of nodes (273,192 compared to 507,822) and a smaller overall size (2.57 versus 2.86 Gb) than our previous pangenome [5]. The core (90%) and flexible (10%) genomes were also slightly lower than those reported in the same study probably due to the exclusion of the X chromosome and a lower number of assemblies (16 vs. 64 individuals). Furthermore, the study by Dai *et al.* [41], which used 20 pseudo-phased HiFi assemblies from Chinese indicine cattle, reports even smaller core genome proportions (78%) and larger flexible regions (22%), reflecting their greater genetic divergence from taurine breeds.

Systematic analysis of the flexible pangenome regions identified 101,219 SVs, of which∼92% were bi-allelic, predominantly insertions and deletions (80%). The remaining SVs, with non-zero reference and alternative allele lengths, were categorized as sequence substitutions.

Comparison with our previous SV panel from 64 assemblies revealed the identification of common 28,488 SVs (28%) with identical coordinates (same start and end positions), and 32,507 SVs (32%) showing at least 90% reciprocal overlap. Size distributions of insertions and deletions were consistent with two previous studies [5][11], showing most variants were small (50 nucleotides to 2 kb) and exhibiting peaks at 150-300bp, 1.5 kb, and 8.5 kb, corresponding to transposable elements. Our results corroborate previous findings which highlighted how transposable elements contribute to structural variation and demonstrated that deletion and insertion events significantly contribute to genomic differences in other species [42][43].

Several SVs detected in our panel are known to be associated with phenotypic traits in cattle. For example, in the Normande breed, we detected an 8.4 kb LINE1 insertion at the ASIP locus on BTA13, which encodes the AGOUTI signalling protein that plays a role in mammalian pigmentation [44]. Additionally, segmental duplications upstream of the *KIT* gene were observed in Abondance, Normande, Montbéliarde, and Simmental breeds, consistent with depigmentation in white-headed breeds [4] [5].

By systematically analysing the SV panel, we further identified 33,634 NRUIs covering a total of 34.4 Mb, of which 14.7 Mb were specific to the yak genome. The remaining 19.7 Mb were identified in at least one *Bos taurus* breed. Approximately 56% of these B. taurus NRUI sequences were shared with our earlier panel, which was constructed using a cattle pangenome developed from 64 de novo assemblies. This led us to combine panels in order to investigate the spectrum of genetic diversity within French cattle breeds. The hierarchical classification based on the genotyping of these NRUIs across a panel of 181 animals from 14 French dairy and beef breeds revealed that all samples were grouped into 14 clusters corresponding to their respective breeds of origin (Fig. 7). These results are of particular interest as they demonstrate the potential of these NRUIs to provide a very clear description of the breed structure. All samples belonging to the same breed are grouped together, without exception. Furthermore, this classification has allowed us to pinpoint four primary clusters of breeds. The first group corresponds to the Holstein breed, which originated in northern Europe and was more recently reintroduced from North America. The second group pertains to the Normande breed, which hails from the Channel area and is significantly influenced by Great Britain. The third group encompasses six breeds (Brown Swiss, Montbéliard, Simmental, Vosgienne, Tarentaise, and Abondance), all of which originated from the eastern

**Figure 7.**
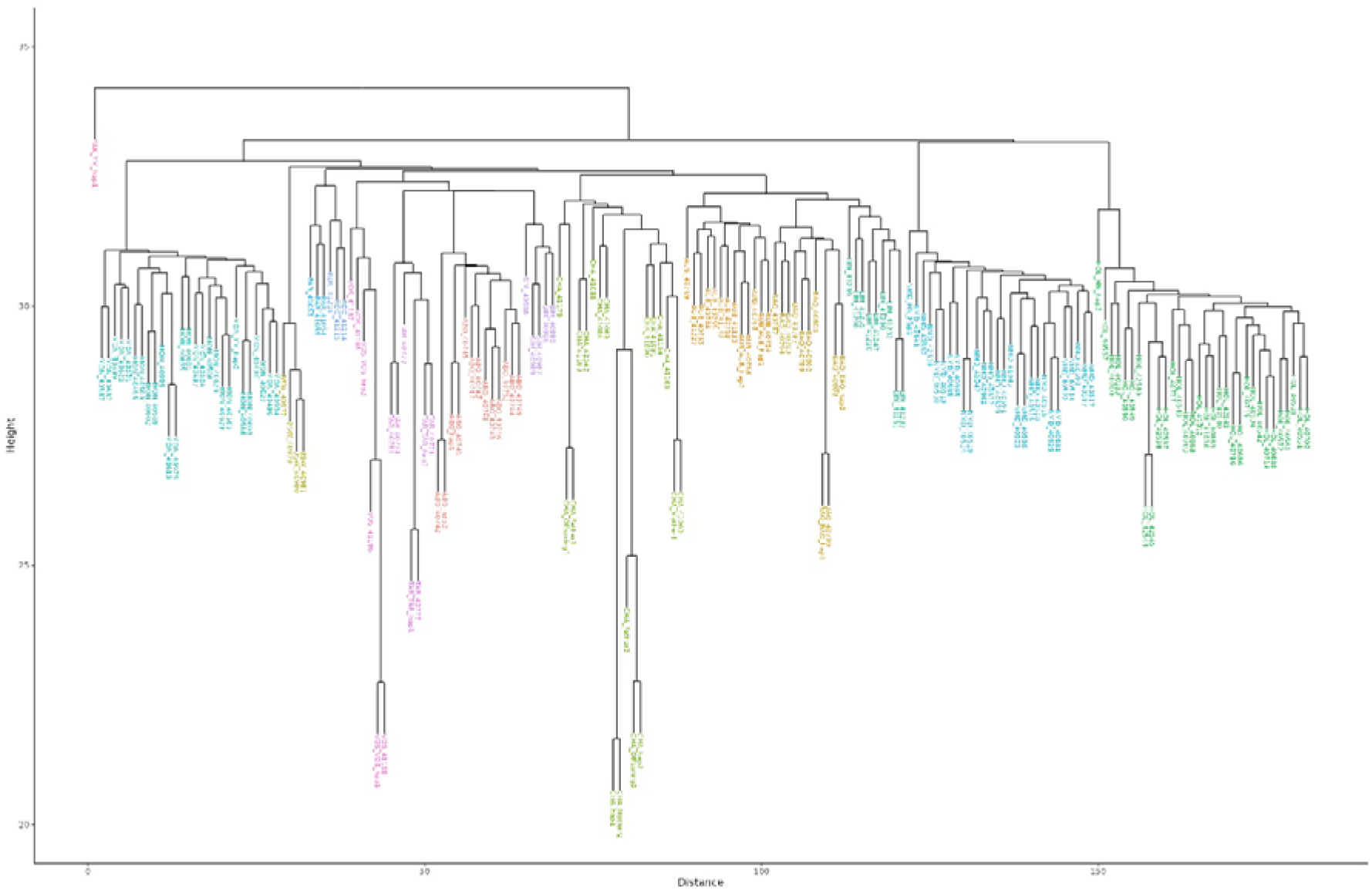
Hierarchical clustering of the 181 *de novo* assemblies based on the NRUI PAV-matrix *PAV* presence/absence variation. Diagram showing the clustering of the 181 assemblies according to the 14 breeds for the 28,998 NRUIs.

Alpine regions. It is important to note, however, that the classification of the Montbéliard breed is further from expected. Lastly, the final group comprises three beef breeds (Limousine, Aubrac, and Blonde d’Aquitaine) that originate from the southwestern region of France and are related to certain Spanish breeds. Additionally, it is worth mentioning that the Charolais breed, which is geographically situated in Central France, is classified between the Alpine and South-West groups, highlighting their dual influence.

Combined annotation efforts along with QTL enrichment and analyses of the distribution of NRUIs between breeds enabled us to pinpoint several NRUIs located within introns of candidate genes. Two Normande-specific NRUIs were identified in *ARMH3* (BTA26:22,672,107; 53 bp, intron 22/25) and *EPHA5* (BTA6:81,086,501; 53 bp, intron 3/16).

The *ARMH3* NRUI significantly increases morphological traits (stature, skeletal development, and chest depth) while reducing milk protein and fat contents without having any effect on milk yields. Although effect on milk yield was not significant, we observed an additive effect of 0.09 GSD (at the heterozygous state) and 0.15 (at the homozygous states). Several studies have already reported significant associations between variants within the *ARMH3* gene and fertility traits in Spring-calved Dairy Cows in New Zealand [45], milk fatty acids in Dual-Purpose Belgian Blue cows [46], and with carcass merit traits in a multibreed population of beef cattle [47]. Also, a genome analysis of the distribution of ROH patterns in Ayrshire cows revealed the identification of several candidate genes, including *ARMH3*, associated with milk production traits, that are under selection pressure [48].

The *ARMH3* gene has been found as implicated in the regulation and structural organization of the Golgi apparatus [49][50]. The Golgi complex plays a central role in protein maturation, lipid processing and vesicular export [51]. Because these functions represent a major role for both milk protein and lipid secretion [52] and skeletal tissue development in animals [53], the Golgi is directly involved in resources allocation between milk production and growth. Consequently, a variation within *ARMH3* gene could modulate the normal function of the Golgi apparatus, therefore influencing the allocation of metabolic resources.

The *EPHA5* NRUI BTA6 at position 81,086,501 exerted a strong influence on milk protein content without affecting milk yield. Since the casein gene cluster is located around 85 Mb on the same chromosome, we assessed the linkage disequilibrium (LD) between the NRUI status and the genotypes of the casein markers included in the EuroGMD genotyping array [54]. This analysis aimed to determine whether the observed effect on the NRUI could be explained by casein genes variants. Across 6 markers in casein genes tested, we observed low to moderate LD values (R^2^ ranging from 0.0002 to 0. 6442, with an average of 0.1864), suggesting that the detected effect most likely originates from the NRUI in *EPHA5*, rather from the casein region. These results are in agreement with previous studies in cattle and sheep, which reported notable correlations between variants in *EPHA5* and both milk and fat production in dairy cattle adapted to tropical climates [55], as well as in fat deposition in domestic sheep breeds originating from Africa and Eurasia [56]. Another study using iHS analysis has pinpointed several genes, including *EPHA5*, currently under selection. This gene is known to play roles in neural development and synaptic functions, possibly contributing to adaptive traits in diverse environments [57].

## Conclusions

This study demonstrates the potential of the pangenome graph approach in comprehensively identifying structural variants, with a particular focus on non-reference unique insertions (NRUIs), providing new insights into genetic diversity within cattle breeds. Our results highlight the potential of pangenomes to enhance reference genomes and uncover genetic variations that are missed using linear assemblies. Importantly, the NRUIs characterized here show strong associations with breed-specific phenotypic traits, providing valuable resources for future research in cattle genetics and breeding programs. It would be very interesting in future work to convert these NRUIs into potentially virtual SNPs, as previously described [58], to add them to the custom EuroGMD SNP chip for genotyping, and to be able to genotype several hundred thousand animals with phenotypes. This data will then be used to explore potential links with phenotypes of interest on a larger scale.

## List of abbreviations

SV: Structural variations
NRUI: Non-reference unique insertions
GWAS: Genome-wide association study
SNP: Single nucleotide polymorphism
GSD: genetic standard deviation
ABO: Abondance
AUB: Aubrac
BAQ: Blonde d’Aquitaine
CHA: Charolais
HOL: Holstein
MON: Montbéliarde
NMD: Normande
TAR: Tarentaise
VOS: Vosgienne

## Declarations

### Ethics approval and consent to participate

Not applicable

### Consent for publication

Not applicable

### Availability of data and materials

The whole genome sequencing raw datasets used in the current study to construct haplotype-resolved assemblies are available in the European Nucleotide Archive (ENA) under projects accession numbers (1) PRJEB64022 and PRJEB101071 for the Abondance, Tarentaise and Vosgienne trios; and (2) PRJEB64023 and PRJEB68296 for Aubrac, Blonde d’Aquitaine, Holstein x Normand and Yak x Montbéliarde Trios.

The new haplotype-resolved phased assemblies constructed in this study and used to build the graph are available at: https://doi.org/10.57745/D4DJYW.

The two Charolais phased assemblies are available at https://doi.org/10.57745/40STPR

All other assemblies used in this study for genotyping are available in the ENA under project accession numbers PRJEB68295 and PRJEB59364.

The pangenome graph, all paths, the SV VCF file and the panel of NRUIs are available at: https://doi.org/10.57745/OXASZE.

### Authors’ contributions

VS, MB, M-PS, GT-K, DB and LD conceived the study; VS conducted the analysis and wrote the original draft; MB, M-PS, GTK and LD contributed to writing the manuscript; CeG, SF and CE collected samples and realized extraction; CeG supervised PCR and sequencing validation of NRUIs; AC and M-MN helped with LR bioinformatics analysis and SV genotyping; M-MN performed hierarchical cluster classification; SF and FB conducted haplotype construction and statistical analyses; CE, JL, CI and CD and DM conceived the experimental design and supervised the technical aspects of the sequencing activity; CK constructed the haplotype-resolved assemblies; All authors reviewed the manuscript; All authors read and approved the final manuscript.

## Acknowledgements

We are grateful to the Genotoul bioinformatics platform Toulouse Occitanie (Bioinfo Genotoul, https://doi.org/10.15454/1.5572369328961167E12) for providing computing and storage resources.

We thank Claire Kuchly and Caroline Vernette for their help to submit the *de novo* genome assembly sequences in the ENA database.

GeT-PlaGe is a member of France Génomique national infrastructure, funded as part of the “Investissements d’Avenir” program managed by the Agence Nationale de la Recherche (contract ANR-10-INBS-09).

Finaly, we would like to thank the Eliance breeding companies that supplied the doses of the sequenced bulls.

## Additional files

**Additional file 1**

Format: Additional file 1.pdf

Title: D-Genies plot for chromosomal alignment concordance between ARS-UCD1.2 *x*-axis and the 16 phased haplotypes on *y*-axis.

Description: **Fig. S1.** Abondance haplotype (A) 1 and (B) 2. **Fig. S2.** Aubrac haplotype (A) 1 and (B) 2. **Fig. S3.** Blonde d’Aquitaine haplotype (A) 1 and (B) 2. **Fig. S4.** Charolais haplotype (A) 1 and (B) 2. **Fig. S5.** Holstein-Normande haplotype (A) 1 and (B) 2. **Fig. S6.** Tarentaise haplotype (A) 1 and (B) 2. **Fig. S7.** Vosgienne haplotype (A) 1 and (B) 2. **Fig. S8.** Yak-Montbéliarde haplotype (A) 1 and (B) 2.

**Additional file 2**

Format: Additional file 2.xlsx

Title: Supplementary tables

Description: **Table S1.** The estimated mash distance of haplotype-resolved assemblies to the reference genome and the order of integrating the assemblies during pangenome construction. **Table S2.** The predicted genotype of each sample to the two NRUIs derived from alignment of LR sequences to the genome followed SV calling. **Table S3.** Effect of the two NRUIs on cattle key traits. **Table S4.** The list of public RNA-Seq data used for expression validation of the novel genes predicted by Augustus. **Table S5.** Significant enrichment of molecular function terms of cattle genes harboring NRUIs using the Panther enrichment system. **Table S6.** Significant enrichment of protein class terms of cattle genes harboring NRUIs using the Panther enrichment system. **Table S7.** Significant enrichment of pathway terms of cattle genes harboring NRUIs using the Panther enrichment system. **Table S8.** Repeats annotated by RepeatMasker. **Table S9.** The list of novel genes predicted by Augustus. **Table S10.** The average TPM estimated from RNA-Seq sequence alignment to the Augustus-predicted novel gene sequences. **Table S11.** The average TPM estimated from RNA-Seq sequence alignment to the 5 genes without detectable homology.

