## Additional file 1 for "Assembly of a pangenome uncovers novel non-reference unique insertion sequences in cattle highlighting their genetic diversity"

A

B

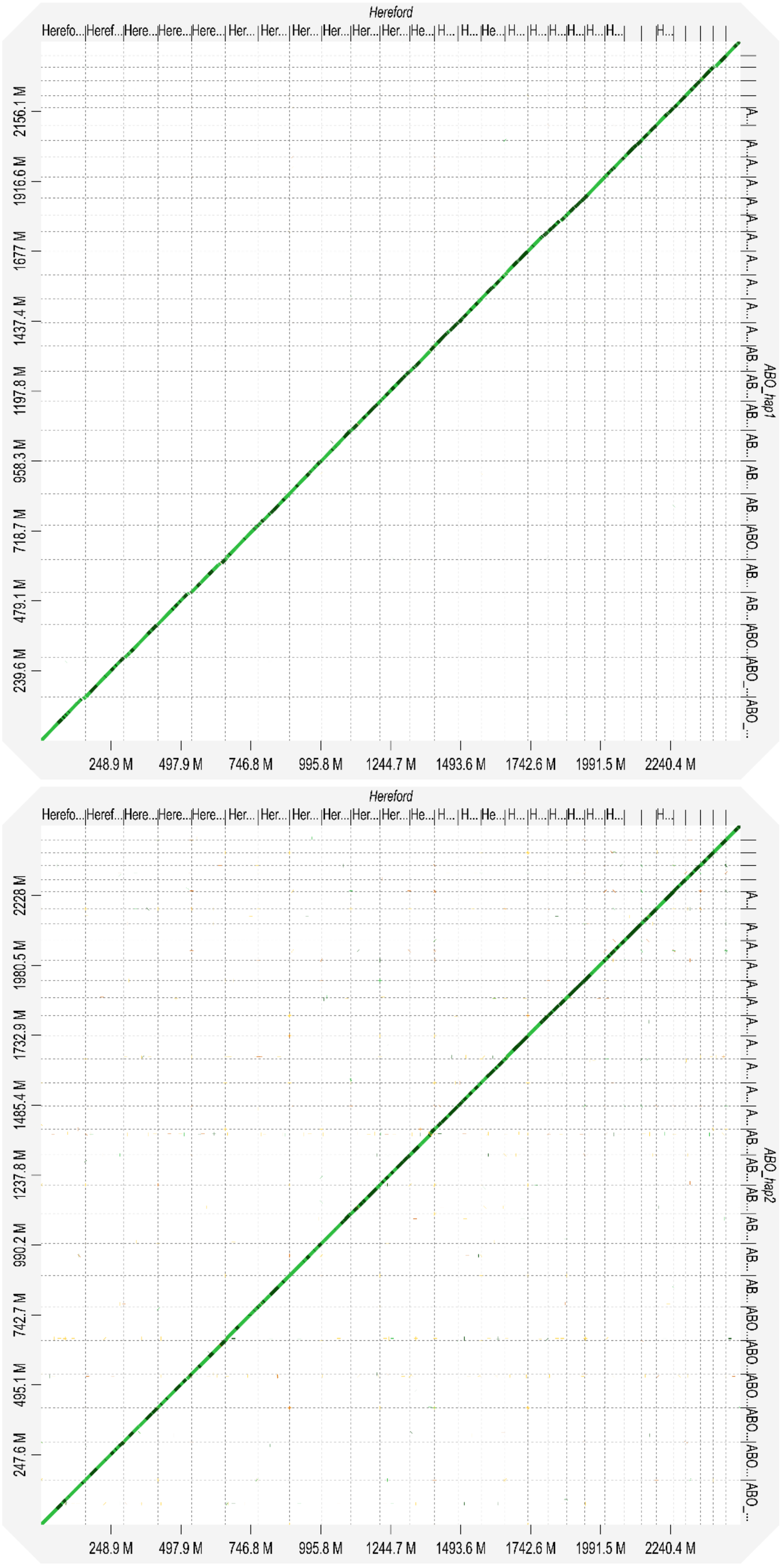


**Fig. S1** D-Genies plot for chromosomal alignment concordance between ARS-UCD1.2 on x-axis and Abondance haplotype (A) 1 and (B) 2 on y-axis.

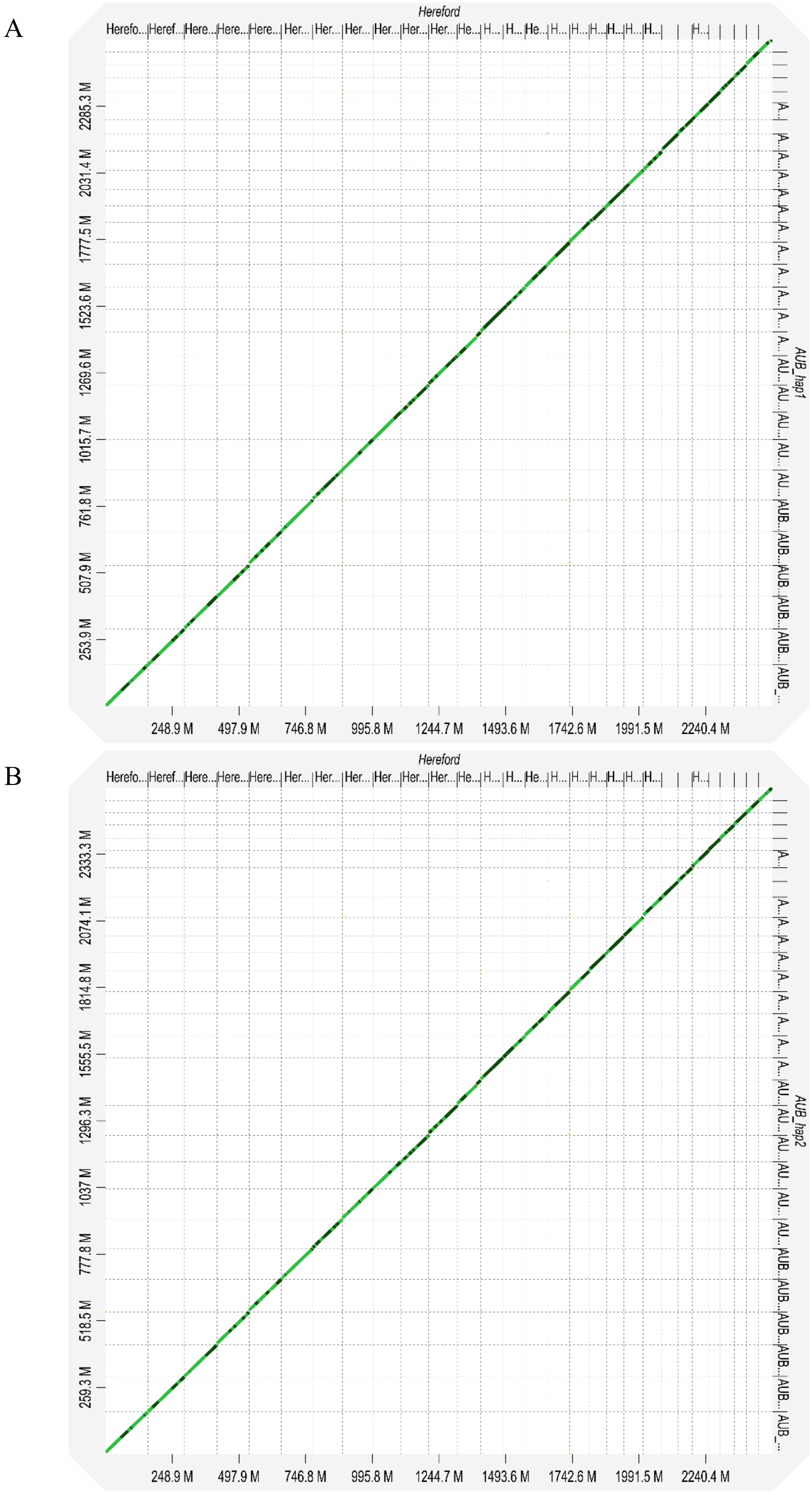


**Fig. S2** D-Genies plot for chromosomal alignment concordance between ARS-UCD1.2 on x-axis and Aubrac haplotype (A) 1 and (B) 2 on y-axis.

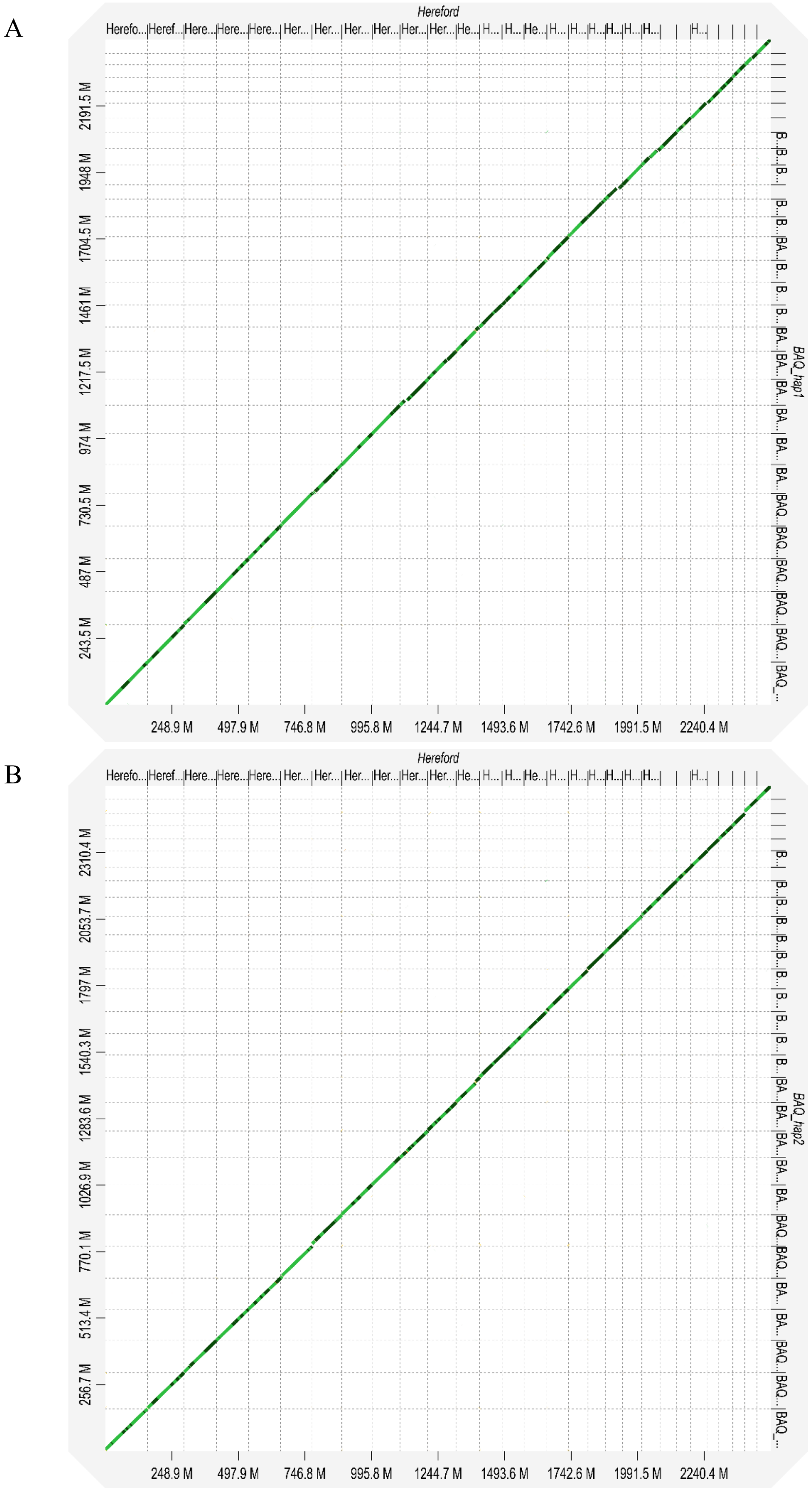


**Fig. S3** D-Genies plot for chromosomal alignment concordance between ARS-UCD1.2 on x-axis and Blonde d'Aquitaine haplotype (A) 1 and (B) 2 on y-axis.

A

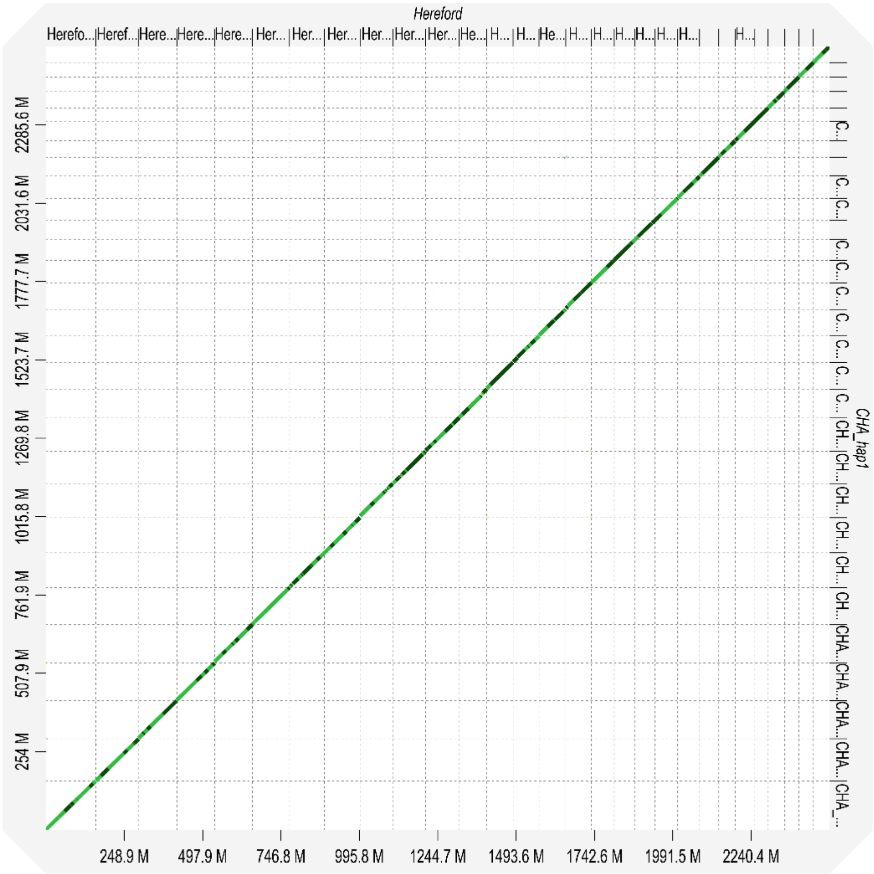


B

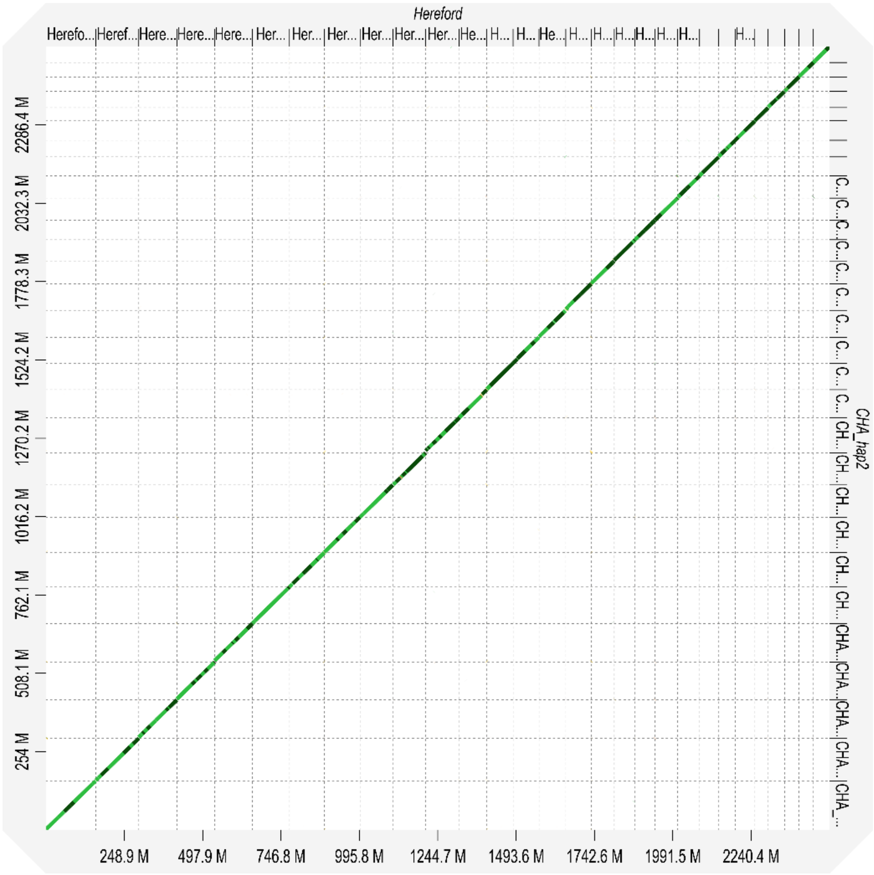


**Fig. S4** D-Genies plot for chromosomal alignment concordance between ARS-UCD1.2 on x-axis and Charolais haplotype (A) 1 and (B) 2 on y-axis.

A

B

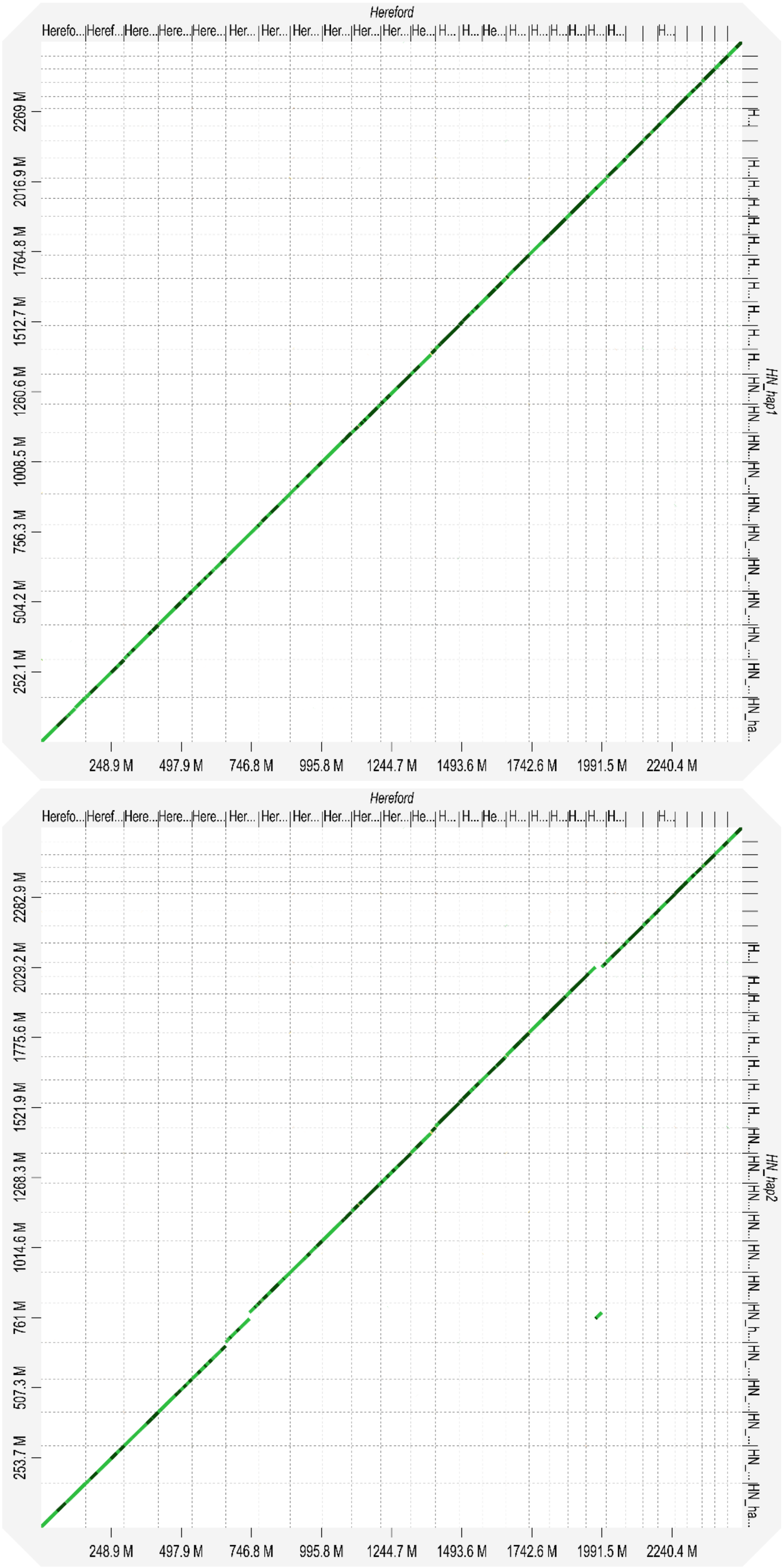


**Fig. S5** D-Genies plot for chromosomal alignment concordance between ARS-UCD1.2 on x-axis and Holstein-Normande haplotype (A) 1 and (B) 2 on y-axis.

A

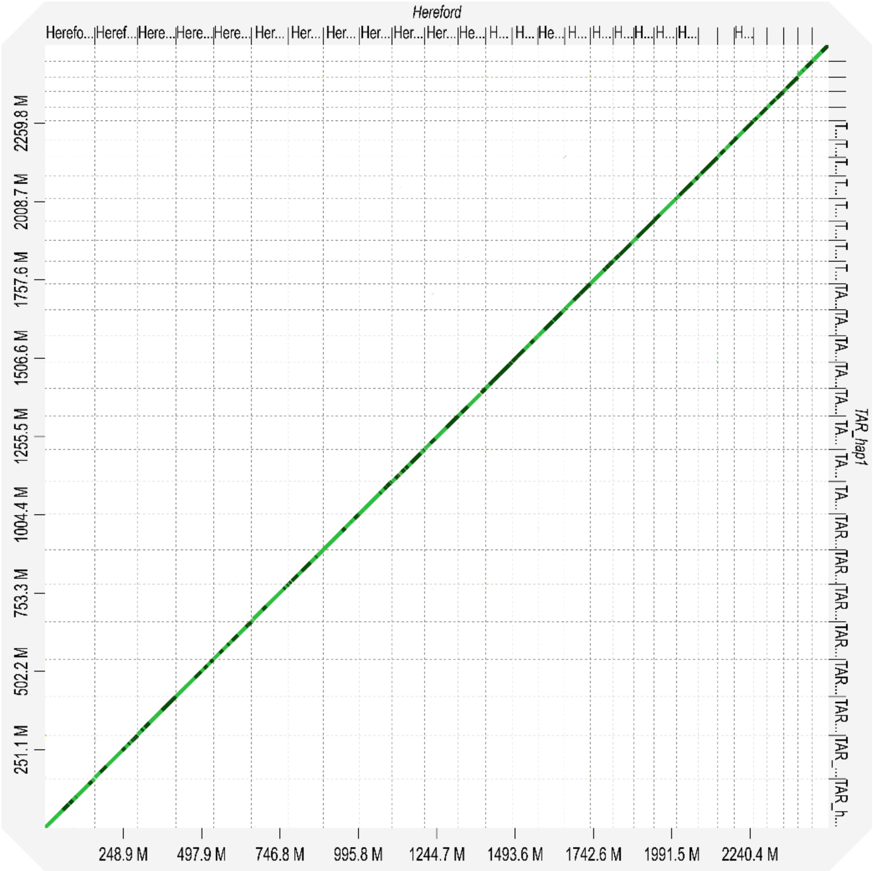


B

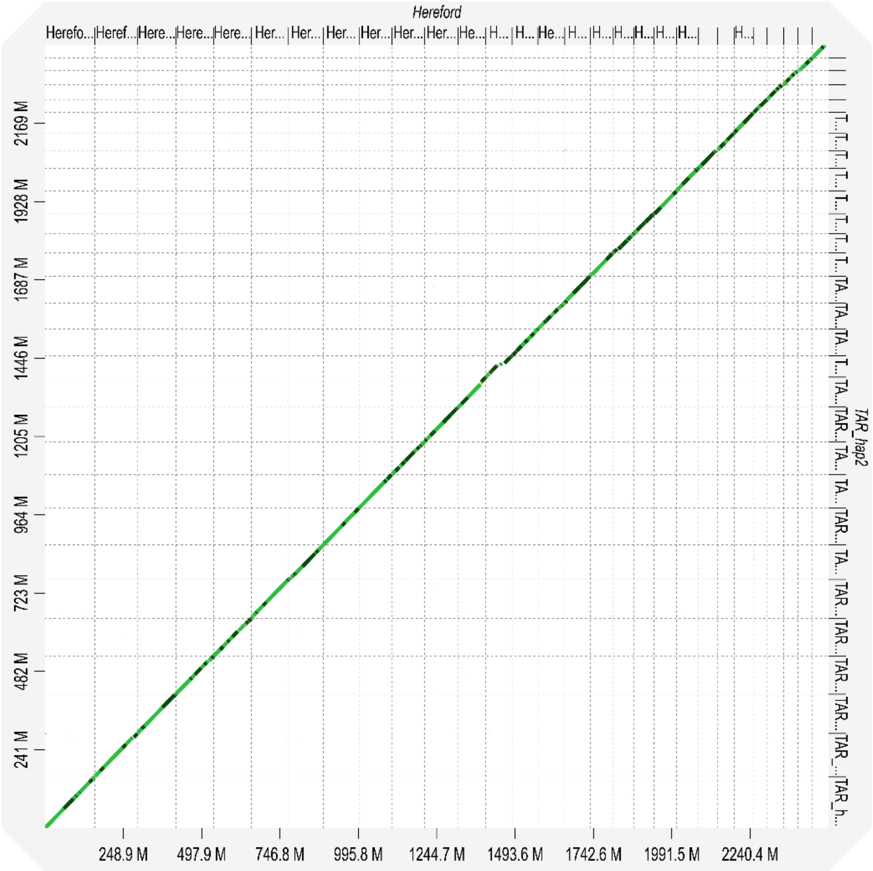


**Fig. S6** D-Genies plot for chromosomal alignment concordance between ARS-UCD1.2 on x-axis and Tarentaise haplotype (A) 1 and (B) 2 on y-axis.

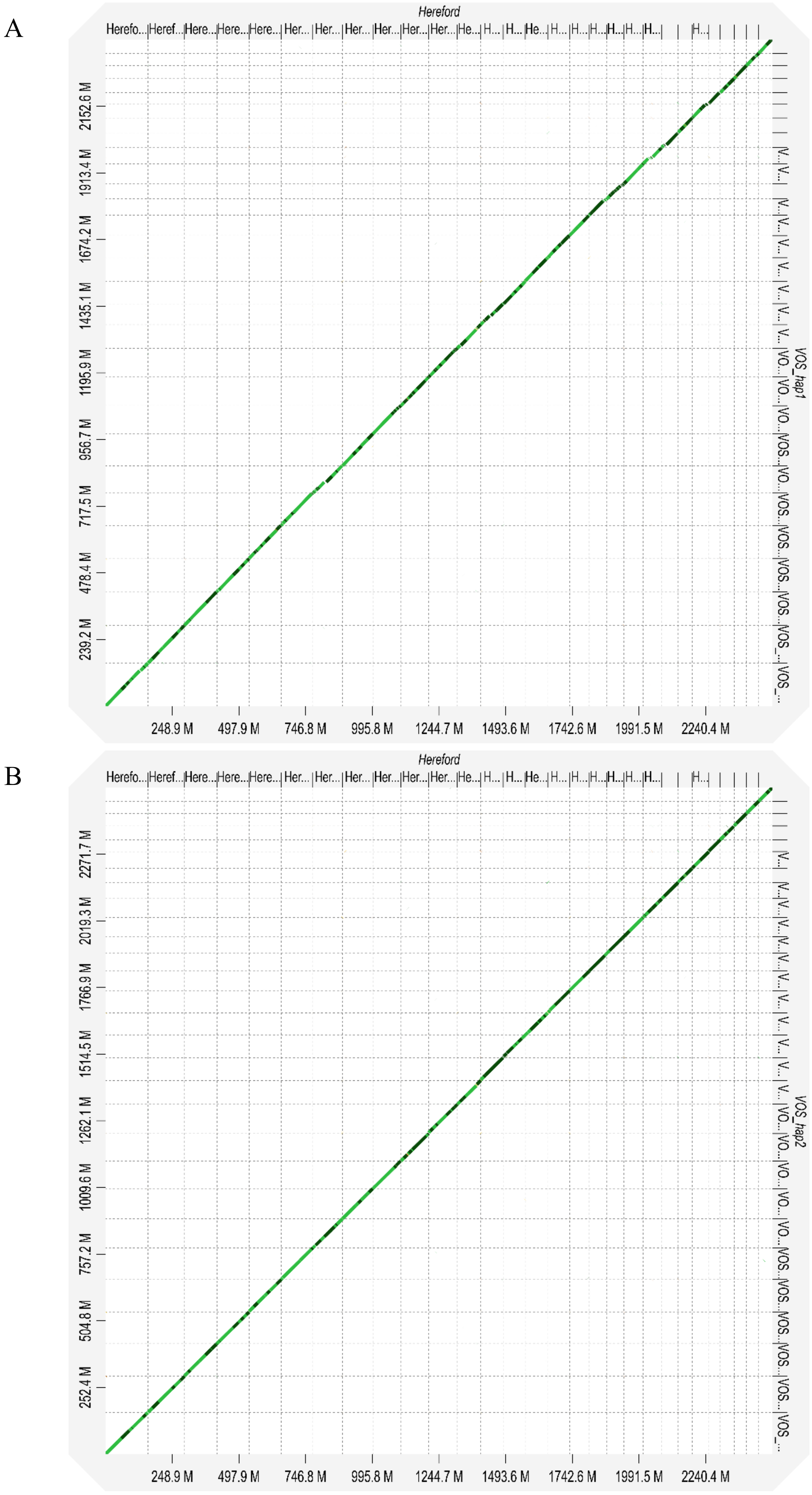


**Fig. S7** D-Genies plot for chromosomal alignment concordance between ARS-UCD1.2 on x-axis and Vosgienne haplotype (A) 1 and (B) 2 on y-axis.

A

B

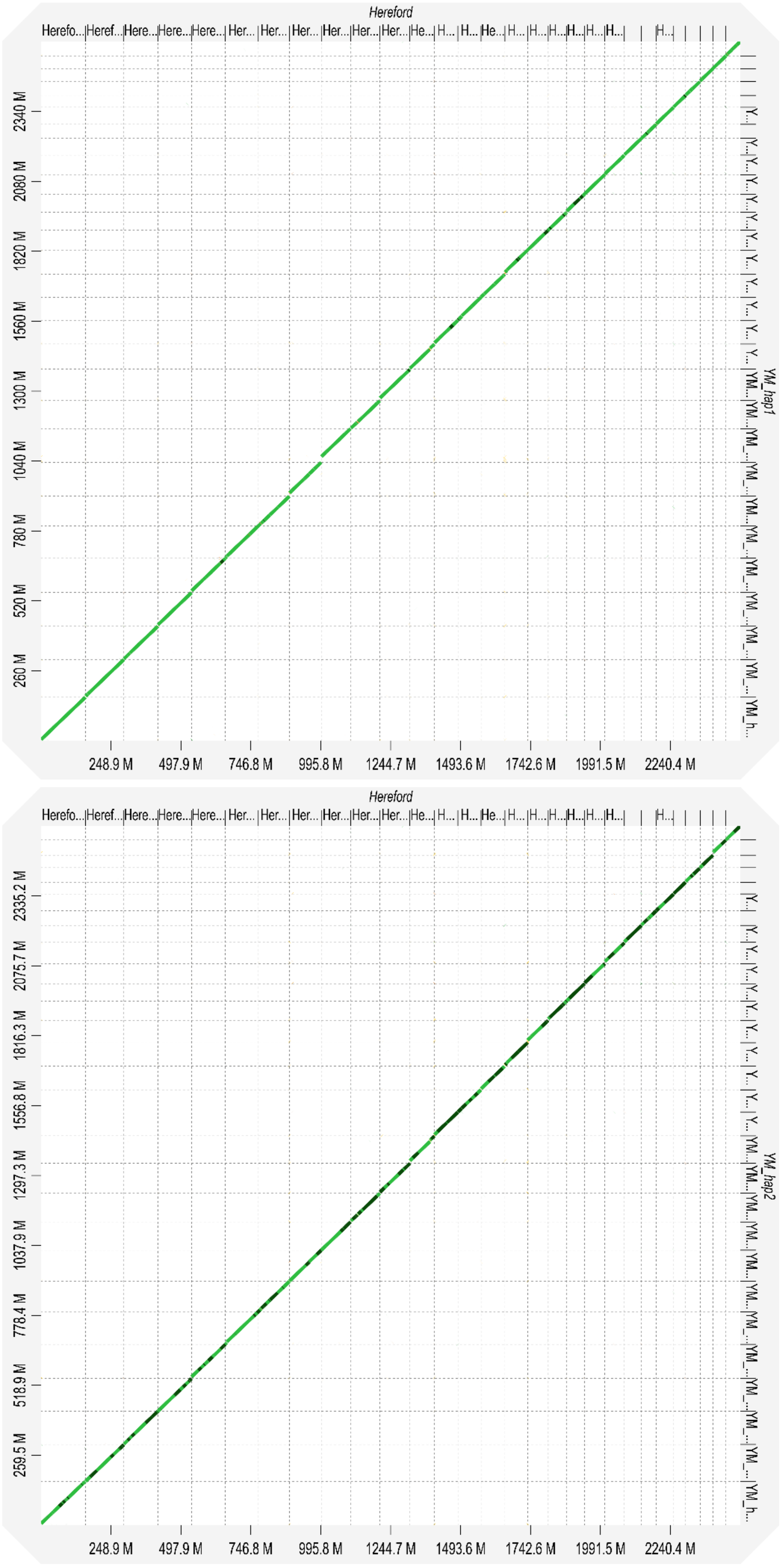


**Fig. S8** D-Genies plot for chromosomal alignment concordance between ARS-UCD1.2 on x-axis and Yak-Montbéliarde haplotype (A) 1 and (B) 2 on y-axis.
